# A butterfly pan-genome reveals a large amount of structural variation underlies the evolution of chromatin accessibility

**DOI:** 10.1101/2022.04.14.488334

**Authors:** Angelo A. Ruggieri, Luca Livraghi, James J. Lewis, Elizabeth Evans, Francesco Cicconardi, Laura Hebberecht, Stephen H. Montgomery, Alfredo Ghezzi, José Arcadio Rodriguez-Martinez, Chris D. Jiggins, W. Owen McMillan, Brian A. Counterman, Riccardo Papa, Steven M. Van Belleghem

## Abstract

Despite insertions and deletions being the most common structural variants (SVs) found across genomes, not much is known about how much these SVs vary within populations and between closely related species, nor their significance in evolution. To address these questions, we characterized the evolution of indel SVs using genome assemblies of three closely related *Heliconius* butterfly species. Over the relatively short evolutionary timescales investigated, up to 18.0% of the genome was composed of indels between two haplotypes of an individual *H. charithonia* butterfly and up to 62.7% included lineage-specific SVs between the genomes of the most distant species (11 Mya). Lineage-specific sequences were mostly characterized as transposable elements (TEs) inserted at random throughout the genome and their overall distribution was similarly affected by linked selection as single nucleotide substitutions. Using chromatin accessibility profiles (i.e., ATAC-seq) of head tissue in caterpillars to identify sequences with potential *cis*-regulatory function, we found that out of the 31,066 identified differences in chromatin accessibility between species, 30.4% were within lineage-specific SVs and 9.4% were characterized as TE insertions. These TE insertions were localized closer to gene transcription start sites than expected at random and were enriched for several transcription factor binding site candidates with known function in neuron development in *Drosophila*. We also identified 24 TE insertions with head-specific chromatin accessibility. Our results show high rates of structural genome evolution that were previously overlooked in comparative genomic studies and suggest a high potential for structural variation to serve as raw material for adaptive evolution.

## Introduction

Structural variants (SVs) in genomes, broadly defined here as encompassing insertions or deletions of at least 1 bp, are a ubiquitous component of within and between species genomic variation (Zhang et al. 2021; Mérot et al. 2020). The larger size of SVs, when compared to single nucleotide polymorphisms (SNPs), may increase their likelihood of being involved in maladaptation (Collins et al. 2020). However, there are a growing number of examples of SV’s role in adaptive innovations (Lucek et al. 2019; Wellenreuther et al. 2019). For example, increased linkage disequilibrium and recombination suppression within large inversions can initiate co-adaptation of gene complexes (e.g., supergenes) in the re-arranged genomic haplotype (Jay et al. 2021; Matschiner et al. 2022). Alternatively, insertion-deletion mutations (indels) can include one or multiple functional genetic elements and studies are starting to indicate that genomic indel content might be large relative to the more commonly studied Single Nucleotide Polymorphisms (SNPs). A study of humans found 2.3 million indels of 1 to 49 bp in length and 107,590 indels larger than 50 bp that accounted for up to 279 Mb in sequence differences among individuals (Ebert et al. 2021). Another case are *Oedothorax* dwarf spiders, in which a large 3 Mb indel is associated with an elaborate alternative reproductive male morph (Hendrickx et al. 2022).

A major challenge in studying the relationships between SVs and adaptive diversification has been the difficulty in characterizing the landscape of divergence in repetitive and rearranged regions of genomes. To overcome this, we here used high-quality butterfly genomes of three *Heliconius* species common to Central and South America and constructed a pan-genome alignment that allowed us to quantify the homologous and non-homologous (i.e., lineage-specific insertions or deletions) portions of their genomes. *Heliconius charithonia* is about 11.1 (8.8-13.4) MY divergent from *H. melpomene* and 6.0 (4.8-7.4) MY divergent from *H. erato* (Kozak et al. 2015; Cicconardi et al. In prep.) (Figure 1A). With the pan-genome alignment, we first analyzed the frequency, length distribution, and composition of lineage-specific sequences between the species. This comparative genome-wide quantification strategy for SVs demonstrated their high abundance is mainly driven by transposable element (TE) accumulation and that SVs can underly more than tenfold sequence differences compared to SNPs between two haploid genomes of a single individual.

**Figure 1.**
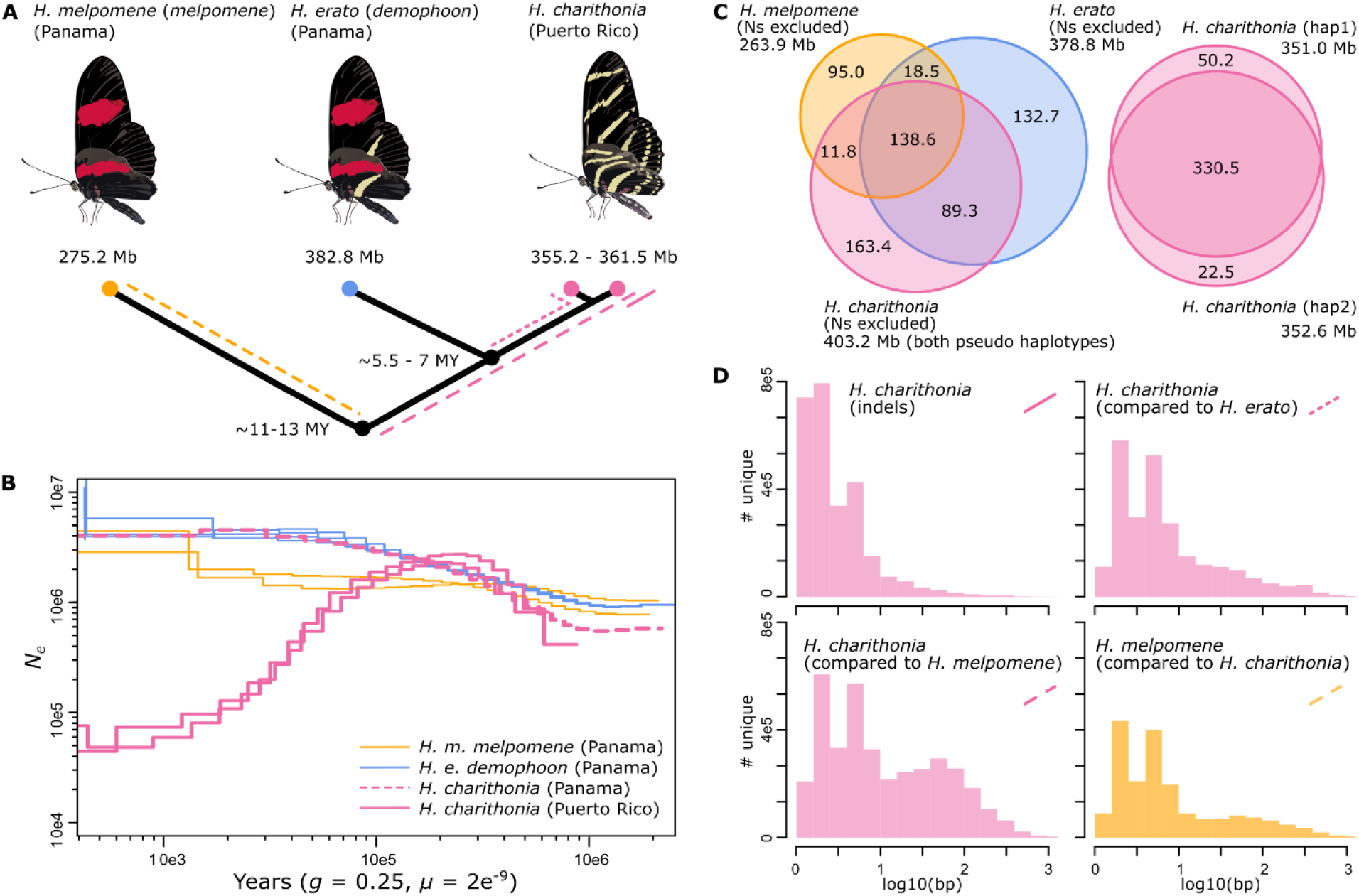
Genome divergence, lineage-specific sequence distribution and historical demography of *H. melpomene, H. erato* and *H. charithonia* from Panama and Puerto Rico. **(A.)** Phylogenetic relations, genome sizes and approximate divergence times. Colored lines indicate branches investigated in panel D. **(B.)** Inference of historical effective population size changes using pairwise sequentially Markovian coalescent (PSMC) analysis. The PSMC estimates are scaled using a generation time of 0.25 years and a mutation rate of 2e-9. Note that the *H. charithonia* genome was obtained from the Puerto Rican population. **(C.)** Venn diagrams represent homologous and non-homologous (lineage-specific) genomic sequences (excluding Ns). Between the two pseudo-haplotypes of the *H. charithonia* genome, we observed a total of 72.7 Mb of sequence identified as indel. Of these indels, 63.1 Mb (86.8%) were lineage-specific to *H. charithonia*, whereas 9.6 Mb (13.2%) were present in the *H. erato* genome. Consistent with divergence times, the *H. charithonia* genome comprised 43.5% (175.2 Mb; compared to the ∼6 MY divergent *H. erato*) to 62.7% (252.3 Mb; compared to the ∼12 MY divergent *H. melpomene*) of lineage-specific sequence resulting from SVs. *Heliconius erato* had 39.0% (151.2 Mb) lineage-specific sequences compared to *H. charithonia* and 58.0% (222.1 Mb) lineage-specific sequences compared to *H. melpomene. Heliconius melpomene* had 34.5% (95.0 Mb) lineage-specific genomic sequence compared to *H. erato* and *H. charithonia*. **(D.)** Length distribution of lineage-specific sequences. Between the two *H. charithonia* haplotypes, indels had an average and median length of 13.5 and 2 bp. The average and median length was 34.2 and 4 bp for lineage-specific *H. charithonia* sequences relative to *H. erato* and 45.6 and 6 bp relative to *H. melpomene*.

Second, we studied the evolutionary processes affecting the distribution and frequency of SVs. We expected that if SVs have a higher chance of being maladaptive, we will see a lower abundance of SVs on smaller chromosomes compared to SNPs. This expectation is derived from smaller chromosomes having a higher per base pair recombination rate that could lead them to purge maladaptive SVs more efficiently (Hill and Robertson 1966). In contrast, if SVs have a similar maladaptive load as SNPs, we expected their abundance on chromosomes to be similar to SNPs, which have a higher abundance on smaller compared to larger chromosomes in *Heliconius* resulting from the higher recombination rate and thus lower reduction of SNP diversity by linked selection on smaller chromosomes (Cicconardi et al. 2021; Martin et al. 2019). To further understand the maladaptive impact of SVs, we also characterized the distribution of SVs relative to gene density. Our hypothesis is that if intergenic SVs impact gene functioning negatively, then we expected to identify fewer SVs in gene rich regions. Moreover, if SVs negatively impact gene regulation, we expected their distances from the transcription start sites (TSS) of genes to be further compared to a random sample of genome positions.

Third, in contrast to maladaptive impacts of SVs, differences in the presence and/or accessibility of *cis*-regulatory loci (i.e., non-coding functional regions of the genome that influence patterns of gene expression) between divergent populations have been shown to house regulatory elements responsible for adaptive differences within and between species of *Heliconius* butterflies (Lewis et al. 2020, 2019; Livraghi et al. 2021). Therefore, to investigate the functional significance of intergenic SVs, we annotated our pan-genome with assays of chromatin accessibility, a powerful approach to identify active *cis*-regulatory sequences (Buenrostro et al. 2013). We focused on chromatin profiles of developing head tissue and wings as a control and observed that lineage-specific open chromatin is substantially associated with SVs. To investigate whether these lineage-specific open chromatin regions within SVs have been involved in recent adaptive evolution, we used selective sweep scans. We also correlated their abundance with gene density and TSS and compared this correlation to that of SVs that do not associate with lineage-specific changes in chromatin accessibility. Finally, using motif enrichment scans for transcription factor (TF) binding sites homologous to *Drosophila*, we investigated whether these lineage-specific SVs carry a high potential for structural variation to serve as material for adaption. In summary, our work here provides a uniquely comprehensive test for the role of SVs in adaptive evolution.

## Results & Discussion

### 1. Genome assemblies, pan-genome alignment, and lineage-specific sequence composition

We *de novo* sequenced and assembled two haploid genomes from a single *H. charithonia* individual from Puerto Rico using 10X chromium technology (10x Genomics, San Francisco, USA). The two pseudo-haploid *H. charithonia* genomes had a length of 355.2 Mb and 361.5 Mb. For *H. erato* and *H. melpomene* we used previously published reference genomes from individuals from Panama which had assembly lengths of 382.8 Mb and 275.2 Mb, respectively (Van Belleghem et al. 2017; Davey et al. 2016). All assemblies had a BUSCO completeness higher than 98.9% (Table S1).

Effective population size influences genetic diversity in SNPs (Charlesworth 2009; Leffler et al. 2012) and is thus also likely to be a major influence on indel diversity. We therefore reconstructed the historical population sizes from diversity estimates from whole-genome resequenced samples using PSMC (pairwise sequentially Markovian coalescent). These reconstructions suggest that populations from Panama have shared a general demographic trend of an increase in population size over the past 1 Mya, with *H. erato* and *H. charithonia* having a larger population size than *H. melpomene* over the last 300 Kya (Figure 1B). In contrast, two *H. charithonia* individuals from Puerto Rico suggest a population size decline over the past 200 Kya. This fits divergence estimates from mtDNA of the Puerto Rican population (Davies and Bermingham 2002) and implies that the indel diversity as estimated from the pseudo-haplotypes of the single Puerto Rican *H. charithonia* individual discussed further on in this study may be a general underestimate of indel diversity in other species or populations such as those from Panama.

For this study, we aligned the four genomes (two *H. charithonia* pseudo-haplotypes, *H. erato* and *H. melpomene*) into a pan-genome with a total length of 659.4 Mb. Among the three species, only 138.6 Mb (21.0%) of sequence was identified as homologous. However, this conserved sequence part retained a high BUSCO completeness of 94.9%, demonstrating it contains the highly conserved gene coding fraction of the genome (Table S1). Such remarkable differences in genome content are also becoming more obvious in other comparative genome studies that incorporate SVs in their analysis, including comparisons between humans and chimpanzee for which genome similarity is much lower than the 99% estimated from the first comparative genomic studies that only considered SNPs and small indels (Mikkelsen et al. 2005; Suntsova and Buzdin 2020). When investigating the proportions of non-homologous (lineage-specific) sequences as obtained from the pan-genome, we found that lineage-specific sequence proportion increases with phylogenetic distance (Figure 1C). More divergent phylogenetic comparisons also had lineage-specific sequences that were generally longer (Figure 1D), whereas less divergent phylogenetic comparisons had a higher proportion of lineage-specific sequences being accounted for by single base pair insertions (e.g., 25.8% of lineage-specific sequence between the *H. charithonia* haplotypes versus 5.76% of lineage-specific sequences between *H. charithonia* and *H. erato*; Table S2).

In the different genome comparisons, we could determine the identity of 61.63 to 70.03% of the lineage-specific sequences (Table S2). TEs constituted the most abundant part (27.0 to 37.0%), which was higher than a previous report that TEs comprise about 26 and 25% of the *H. erato* and *H. melpomene* genome sequence, respectively (Lavoie et al. 2013; Ray et al. 2019). While insertions, deletions, duplications, and inversions can all cause SVs, transposable elements (TEs) are thus identified as the most common source of SVs, similar to many other organisms (Garcia-Perez et al. 2016; Cerbin and Jiang 2018).

Among phylogenetic comparisons, we found generally similar patterns of TE family accumulation but observed several lineage-specific differences (Figure 2). The most abundant elements associated with lineage-specific sequences in all genome comparisons were SINE elements (25 to 41%), Rolling-circle elements (23 to 35%), LINE elements (10.1 to 22.4%), and DNA transposable elements (14.8 and 21.25%) (Figure 2A). Between the two *H. charithonia* haplotypes the two most abundant groups associated with indels were Rolling-circle (32.5%) and SINE (29.7%), with Helitron2_Hera and Metulj7_Hmel showing highest copy numbers (6.5 and 3.6% variation in activity, respectively; Figure 2B).

**Figure 2.**
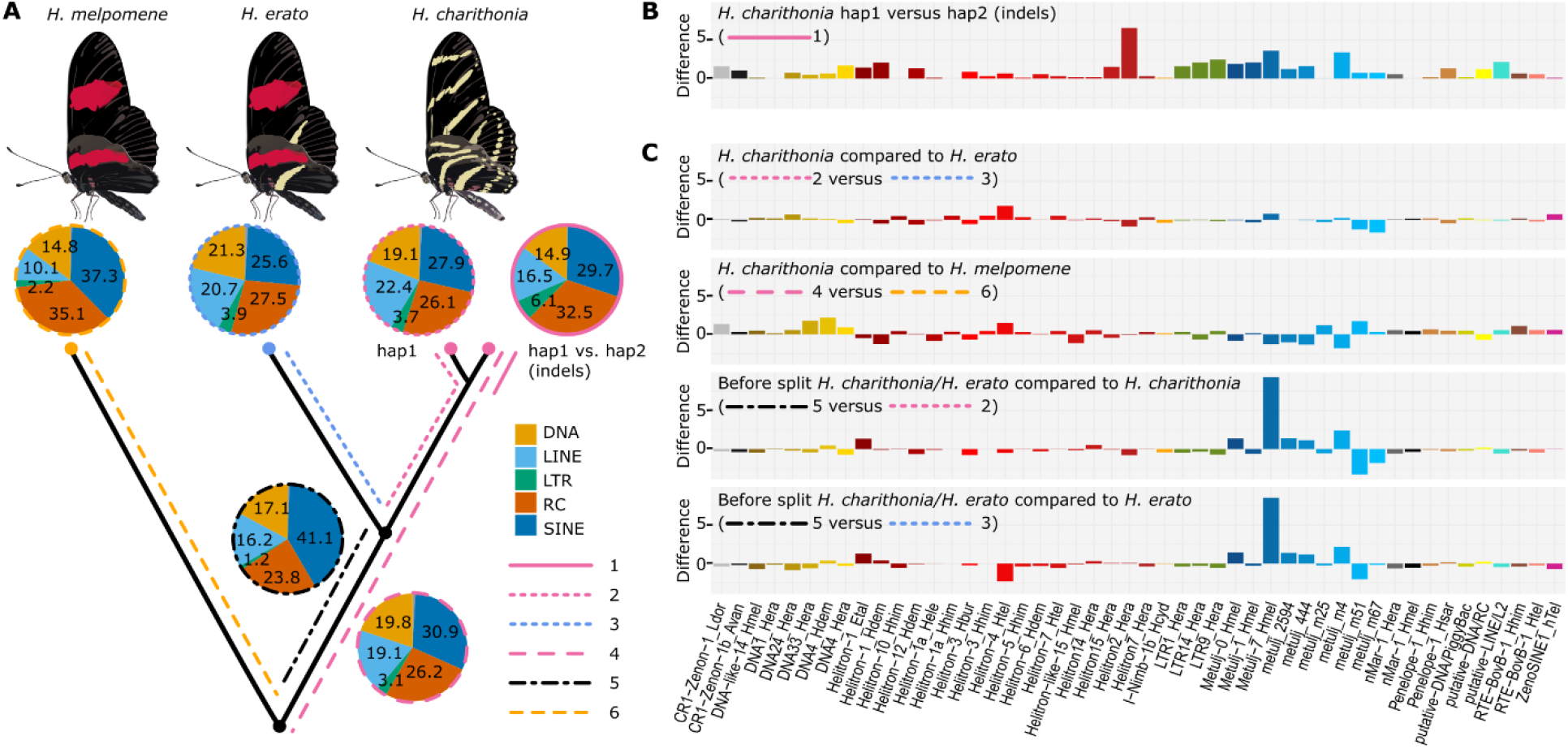
Phylogenetic dynamic of Transposable Elements (TEs). **(A)** Lineage-specific TE family accumulation. Different line types depict different branches in the phylogeny studied and allow to investigate changes in temporal accumulation of TEs. 1 = TE families associated with indels between the *H. charithonia* haplotypes, 2 = TE families accumulated in *H. charithonia* since the split from *H. erato*, 3 = TE families accumulated in *H. erato* since the split from *H. charithonia*, 4 = TE families accumulated in the *H. erato* lineage since their split from a common ancestor with *H. melpomene*, 5 = TE families accumulated after the *H. charithonia/H. erato* lineage split from the common ancestor with *H. melpomene* but before *H. erato* and *H. charithonia* split, 6= TE families accumulated in the *H. charithonia/H. erato* lineage since their split from a common ancestor with *H. melpomene*. DNA = DNA transposons that do not involve an RNA intermediate; LINE = long interspersed nuclear elements, which encode reverse transcriptase but lack LTRs; LTR = long terminal repeats, which encode reverse transcriptase; RC = transpose by rolling-circle replication via a single-stranded DNA intermediate (Helitrons); SINE = short interspersed nuclear elements that do not encode reverse transcriptase. (**B**.) Difference in TEs (percentage of total) between the two *H. charithonia* haplotypes considering the same 48 most significantly divergent TE families. **(C.)** Difference in TEs (percentage of total) between branches in the phylogeny considering the same 48 most significantly divergent TE families. Positive values indicate higher accumulation in the first branch, negative values indicated higher accumulation in the second branch of the comparison. Total TE accumulation patterns per lineage are shown in Figure S1.

Our phylogenetic framework next allowed us to characterize the time of accumulation for TEs along the *H. erato/H. charithonia* branch (considering *H. melpomene* as the outgroup). Within the TE families, we found that Metulj-7 elements accumulated before *H. erato* and *H. charithonia* split (Figure 2B). This was also supported by relative age of accumulation analysis based on divergence of Metulj-7 that showed accumulation was more ancient than, for example, metulj_m51 that likely increased in number after *H. charithonia* and *H. erato* split (Figure S1). Metulj-7 also accrued earlier in the *H. melpomene* lineage (Figure S1). This may imply an accumulation that precedes the split of our butterfly lineages. The reduction of Metulj-7 in more recent times supports a similar finding by Ray *et al*. (2019), who observed a reduction of Metulj-7 accumulation in the *H. charithonia/erato* lineage starting at 5 Mya (Ray et al 2019).

### 2. Indel patterns and chromosome sizes

We examined if the abundance of SVs across the genome is similarly affected by linked selection as SNP diversity. In *Heliconius*, there is a negative relationship between average nucleotide diversity (i.e., average pairwise nucleotide differences) and chromosome size, with larger chromosomes generally carrying lower diversity (Figure 3A; Martin et al. 2019; Cicconardi et al. 2021). In the case of nucleotide diversity and chromosome size, this negative relationship has been explained by an increased reduction of genetic diversity at linked sites by greater background selection and genetic hitchhiking on larger chromosomes (Cicconardi et al. 2021; Campos and Charlesworth 2019; Cutter and Payseur 2013). Genetic linkage map suggests that there is on average a single crossover per meiosis, regardless of chromosomal length (Davey et al., 2016). This results in longer chromosomes having a lower per-base recombination rate, which increases the extent of linked selection and results in lower nucleotide diversity on larger chromosomes. However, if SVs have a higher maladaptive mutation load because of their size, we might expect the opposite pattern in which shorter chromosomes with higher recombination rates were able to purge SVs more easily through recombination (Hill and Robertson 1966). Thus, there might be a positive relationship between SV diversity and chromosome length.

**Figure 3.**
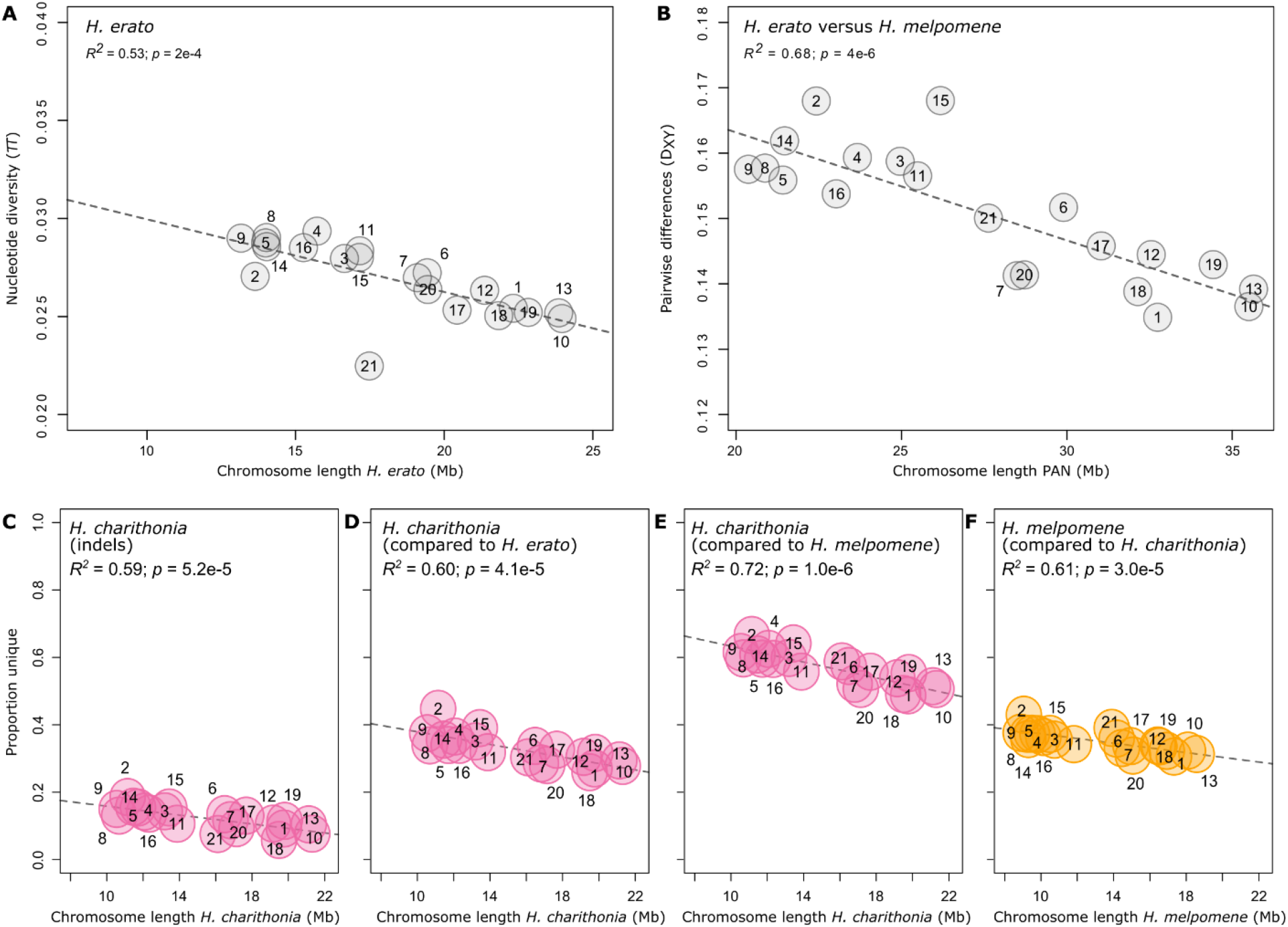
Patterns of lineage-specific sequence distribution and chromosome lengths. **(A.)** Correlation between *H. erato* chromosome lengths and nucleotide diversity (obtained from whole genome resequence data of ten *H. e. demophoon* samples from Panama). **(B.)** Correlation between pan-genome chromosome lengths and pairwise nucleotide differences (*D*_*XY*_) averaged for each chromosome between *H. erato* and *H. melpomene. D*_*XY*_ was calculated from homologous sequences in the pan-genome. **(C.)** Correlation between chromosome lengths and proportion of indels in the chromosomes of *H. charithonia*. Correlation between chromosome lengths and proportion of lineage-specific sequences in the chromosomes of **(D.)** *H. charithonia* compared to *H. erato*, **(E.)** *H. charithonia* compared to *H. melpomene*, and **(F.)** *H. melpomene* compared to *H. charithonia*. Dashed lines indicate regression fit. Numbers indicate chromosome numbers. Colors refer to sequences specific to *H. charithonia* (pink) and *H. melpomene* (orange).

In fact, our data are most consistent with the hypothesis that SVs are affected by linked selection in a manner similar to SNPs. Indeed, between the two pseudo-haplotypes of *H. charithonia*, there was a significant negative relationship between the proportion of indels in each chromosome and chromosome sizes (Figure 3C). This suggests that the general diversity of SVs in a population may be driven by linked selection similar to SNPs. Patterns of the proportion of lineage-specific sequences may then have been largely driven by patterns of ancestral diversity, resulting in higher proportions of lineage-specific sequences on smaller chromosomes (Figure 3D-F), as is also observed for pairwise nucleotide divergence patterns between, for example, *H. erato* and *H. melpomene* (Figure 3B; Van Belleghem et al. 2018). This expectation is borne out on the sex (Z) chromosome (21), where there was a reduction in SV diversity that roughly mirrored the patterns of SNP diversity. Due to its hemizygous state in females, there is a smaller effective population size (0.75 relative to autosomes) and an expected reduction in SNP diversity (Charlesworth 2001). For indels within *H. charithonia*, we found a 0.61 ratio of indel proportion on chromosome 21 compared to the autosomes, suggesting that indels are subject to differences in effective population size similarly to SNPs.

We next characterized the distribution of SVs relative to genes to further explore the potential maladaptive impact of SVs. TEs, the most abundant SVs, are argued to most often have a neutral or negative impact and end up silenced by genome defense mechanisms (Okamoto and Hirochika 2001; Rigal and Mathieu 2011). If intergenic TEs impact gene functioning negatively, we expected to identify fewer TEs in gene rich regions. Moreover, if TEs negatively impact gene regulation, we expected their distances from the 5’-end of genes (as a proxy for the transcription start site (TSS)) to be further compared to a random sample of genome positions. In agreement with the former expectation, the frequency of lineage-specific TEs correlated negatively with gene frequency (*R*^*2*^ = -0.27, *p* < 0.001; Figure 4A), suggesting a general purifying selection against SVs and TEs in gene dense regions. The distance distribution of TEs to TSS was significantly higher than random expectations although visually similar (Figure 4B), which may reflect their tendency to randomly insert in the genome in terms of genomic position.

**Figure 4.**
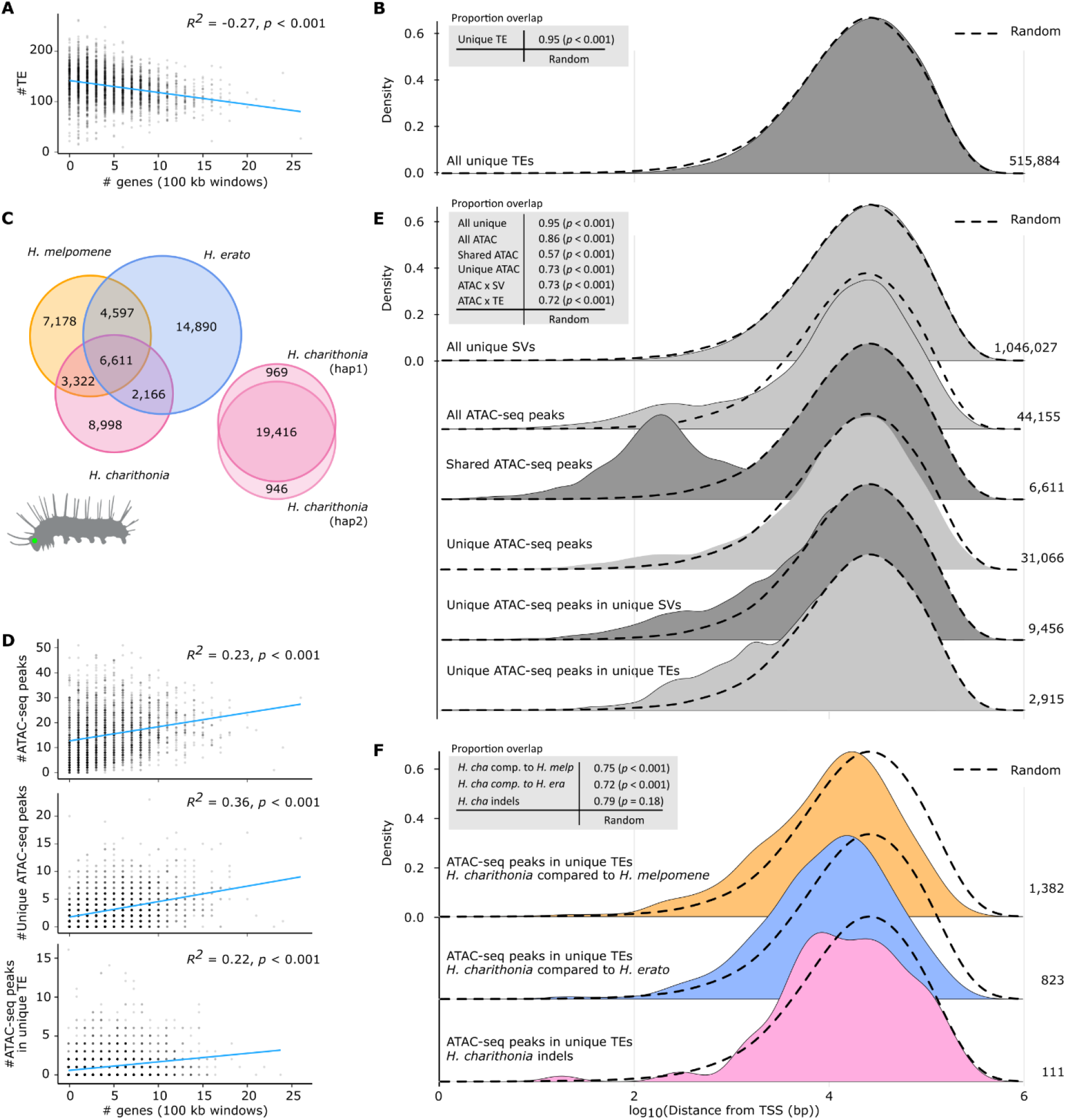
Lineage-specific sequences and their relationship with chromatin accessibility and gene distribution. **(A.)** Correlation of gene density in 100 kb windows with frequency of transposable elements (TEs). **(B.)** Density plot of distance of lineage-specific TEs to closest transcription start site (TSS) pooled over all species genome comparisons. **(C.)** Lineage-specific and shared open chromatin signals (ATAC-seq peaks) found in head tissue of 5th instar caterpillars in each species. Peaks are considered shared (homologous) when they overlap at least 50% reciprocally. **(D.)** Correlation of gene frequency in 100 kb windows with frequency of all lineage-specific structural variants (SVs), all ATAC-seq peaks, lineage-specific ATAC-seq peaks, and lineage-specific TE insertions with ATAC-seq peaks. **(E.)** Density plot of distance of lineage-specific sequence features to closest transcription start site (TSS) pooled over all species genome comparisons. We found the distribution of lineage-specific SVs was most similar to a random distribution of positions in the genome (overlapping index = 95%), with a median/mean distance of 21,701/40,790 bp of a lineage-specific sequence and 20,801/39,908 bp of any random position to a TSS. **(F.)** Density plot of distance of lineage-specific TEs with ATAC-seq peaks in *H. charithonia* to closest TSS. Dashed lines show the distance distribution to TSS of 100,000 randomly selected positions. Tables at the top left in panels B, E and F report overlapping indexes and pairwise Wilcoxon test p-values between the distributions of lineage-specific sequence features and the random positions. Numbers on the right indicate the number of the respective sequence features.

### 3. Genomic landscape of DNA accessibility and functional potential of TEs

Although the genome-wide distribution patterns of SVs and TEs seems to resemble that of SNPs, we next wanted to investigate the functional and adaptive significance of lineage-specific intergenic SVs. TEs, for example, have been suggested to be important genomic material for *cis*-regulatory element evolution (Branco and Chuong 2020; Pontis et al. 2019; Fueyo et al. 2022). To test this, we studied the genomic distribution of potential *cis*-regulatory elements (CREs) using Assays for Transposase-Accessible Chromatin using sequencing (ATAC-seq) (Buenrostro et al. 2013). We obtained ATAC-seq data for head tissue from 5^th^ instar caterpillars, a tissue labile to adaptive change (Montgomery and Merrill 2017; Montgomery et al. 2021) and a developmental stage that can be confidently timed to minimize species differences in developmental rates or heterochrony (Reed et al. 2007). In *H. melpomene, H. erato* and *H. charithonia*, we counted, respectively, 21,708, 28,264 and 21,097 ATAC-seq peaks that significantly represented open chromatin (Figure 4C). Of these peaks, 6,611 (13.8%) of the total recorded peaks were identified as homologous (overlapped at least 50% reciprocally between all three species), whereas 31,066 were lineage-specific.

If open chromatin indeed correlates with active gene regulation, we expected to find more ATAC-seq peaks in gene dense regions of the genome. In agreement with such active gene regulation, ATAC-seq peaks were indeed enriched in regions of the genome with higher gene density (*R*^*2*^ = 0.23, *p* < 0.001; Figure 4D). This positive correlation with gene density was also observed for ATAC-seq peaks that were lineage-specific (*R*^*2*^ = 0.36, *p* < 0.001), and ATAC-seq peaks that were within lineage-specific SVs and TEs (*R*^*2*^ = 0.22, *p* < 0.001), which supports that they may also have *cis*-regulatory activity. Moreover, these ATAC-seq peaks were closer to TSS than random (Figure 4E). Although some of the lineage-specific chromatin accessible peaks may result from differences in developmental timing between the three species, out of these 31,066 lineage-specific peaks, 9,456 (30.4%) were within SVs of which 2,915 (9.4%) could be annotated as TEs. We also note that the absolute number of functional elements within SVs and TEs may be much higher than what is described in our study because we restricted our chromatin data to only one tissue type and developmental time point. As a comparison, a genomic study across 20 mammalian genomes spanning 180 MY of evolution identified roughly half of all active liver enhancers specific to each species, but argued that most of these lineage-specific enhancers evolved through redeployment of ancestral DNA and that a significant contribution of repeat elements to enhancer evolution was only found for more recently evolved enhancers less than 40 Mya old (Villar et al. 2015).

Considering that ATAC-seq peaks identified in the head tissue could also be accessible in the germline, we next investigated if the distribution of lineage-specific ATAC-seq peaks within TEs closer to TSS may have been caused by inserting in open chromatin, or whether these TE insertions may have caused the open chromatin and have been selectively retained at these positions. For this analysis, we looked for chromatin signals in homologous sequences flanking the TEs in the other species. We found that 395 (13.5%) out of 2,915 lineage-specific TEs with ATAC-seq peaks had a significant ATAC-seq signal in the other species within 2,000 bp of homologous sequence flanking the insert. This was higher than an expected 2% obtained from 1,000 random permutations of an equal number of TEs that did not associate with ATAC-seq peaks. Nevertheless, 2,520 (86.4%) did not have any ATAC-seq signal in the other species. To further test whether these TEs have been selectively retained closer to TSS, we performed a TSS distance distribution comparison of SVs within ATAC-seq peaks specific to *H. charithonia* (Figure 4F). A comparison relative to *H. erato* and *H. melpomene* showed significantly closer TSS distances of lineage-specific sequences with ATAC-seq peaks compared to random (Wilcoxon *p*-value < 0.001), whereas the distribution of indels with ATAC-seq peaks within the single *H. charithonia* individual was not statistically different compared to a random distribution of positions in the genome (Wilcoxon *p*-value = 0.18). Therefore, the pattern of TE insertions associated with open chromatin being closer to TSS could have potentially arisen over time because of TE insertions closer to TSS having a higher chance of affecting gene expression and being involved in adaptive changes between species.

Although the distribution of ATAC-seq peaks within TEs can fit selective retention of these SVs, we wanted to directly test for the influence of selection using selective sweep analysis. Given the demographic history of our taxa and using an effective population size of 2 million individuals (Moest et al. 2020), it is important to recognize that our ability to identify signals of adaptation is restricted to selection acting within the past 80,000 years (0.6% of the studied evolutionary timescale). Under these restricted conditions, we did not find a pattern of recent adaptive evolution (Figure S3). Nevertheless, we did observe that TE insertions associated with open chromatin were more fragmented compared to other TEs in the genome, suggesting selection for immobilization of these TEs (Joly-Lopez and Bureau 2018) (Figure S2). Thus, strong signals for recent selection were not obvious in our comparative data.

Nonetheless, even if SVs are mostly neutral or deleterious, their shear abundance and association with chromatin accessibility changes between species underscores their adaptive potential. In agreement with the importance of TEs to adaptive evolution, several examples come from the genomes of Lepidoptera. In the classic example of industrial melanism of the peppered moth, a novel 21 kb TE insertion that impacts function of the gene *cortex* is responsible for the development of the different color morphs (Van’t Hof 2016). Another TE insertion has been linked to the silencing of a *cortex* regulatory region and may be responsible for the yellow band on the hindwing in geographic variants of *Heliconius melpomene* butterflies (Livraghi et al. 2021).

In several studies, TEs have been correlated to evolutionary changes in chromatin state, gene expression, and adaptive evolution at a genome-wide scale (Bourque et al. 2018; Diehl et al. 2020; Ohtani and Iwasaki 2021; Liu et al. 2019). The TE family composition of TEs that associated with open ATAC-seq peaks was markedly different between the three *Heliconius* species (Figure S4). To infer the evolutionary potential of the accumulation of these TE families, we next identified enrichment of sequence motifs in chromatin changes that are within lineage-specific TE insertions and investigated their potential as transcription factor (TF) binding sites. TF binding motif enrichment analysis on the 2,915 lineage-specific ATAC-seq peaks within TE insertions showed that each genome has unique signals of binding site enrichment that were homologous to binding sites of TFs in *Drosophila* (Figure 5). Eight of the identified 21 enriched binding motifs were for TFs with known functions in nervous system development in *Drosophila*. For example, *lola, Dref*, and *shn* are involved in dendrite morphogenesis (Iyer et al. 2013), *Hr51* is related to neuron remodeling and circadian rhythm regulation (Kozlov et al. 2017), *slp2* regulates a wide variety of developmental processes in *Drosophila*, including embryonic segmentation, ventral fate specification in the retina, and temporal patterning of the neuroblasts that produce medulla neurons (Sato and Tomlinson 2007), *wor* regulates neurogenesis (Ashraf et al. 2004), *esg* is involved in central nervous system development (Ashraf et al. 1999), *Btd* contributes to embryonic head segmentation (Wimmer et al. 2010), and *Fer1* is involved in the maintenance of neuronal identity (Guo et al. 2019). Three other TF motifs have been previously linked to wing or color pattern development in lepidoptera. *Mad* is a TF linked to wing development in *H. melpomene* (Baxter et al. 2010). *Mitf* has been associated with color pattern development in other animals (Mallarino et al. 2016; Poelstra et al. 2015), and in *Heliconius* butterflies potentially interacts with *aristaless* (Westerman et al. 2018). Finally, *dsx* controls sex-limited mimicry patterns in *Papilio polytes* and *Zerene cesonia* butterflies (Rodriguez-Caro et al. 2021; Nishikawa et al. 2015).

**Figure 5.**
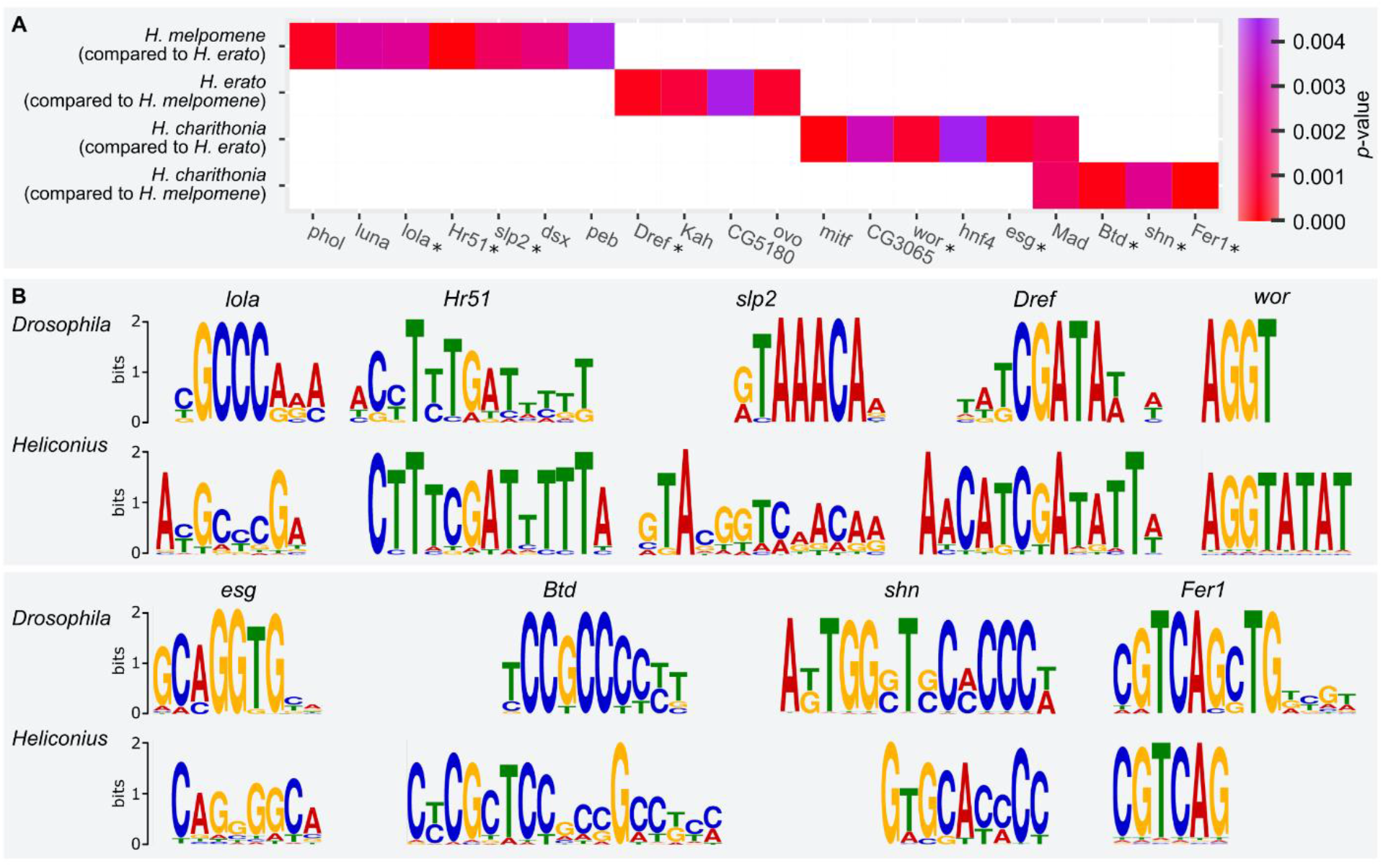
Transcription factor (TF) binding motif enrichment among lineage-specific TE insertions with ATAC-seq peaks. **(A.)** Lineage-specific ATAC-seq peaks within TEs in *Heliconius melpomene* (compared to *H. erato*) were significantly enriched for binding sites with significant homology to flybase *D. melanogaster* TF binding of *pleiohomeotic like* (*phol*; FBgn0035997), *luna* (*luna*; FBgn0040765), *longitudinals lacking* (*lola*; FBgn0283521), *Hormone receptor 51* (*Hr51*; FBgn0034012), *sloppy paired 2* (*slp2*; FBgn0004567), *doublesex* (*dsx*; FBgn0000504), and *pebbled* (*peb*; FBgn0003053). *Heliconius erato* (compared to *H. melpomene*) was significantly enriched for binding sites for DNA replication-related element factor (*Dref*; FBgn0015664), *Kahuli* (*Kah*; FBgn0035144), *CG5180* (*CG5180*; FBgn0043457), and *ovo* (*ovo*; FBgn0003028 *H. charithonia* (compared to *H. erato*) was enriched in binding sites for *Mitf* (*Mitf*; FBgn0263112), *CG3065* (*CG3065*; FBgn0034946), *worniu* (*wor*; FBgn0001983), *hnf4* (*hnf4*; FBgn0004914), *escargot* (*esg*; FBgn0287768), and *Mothers against dpp* (*Mad*; FBgn0011648). *H. charithonia* (compared to *H. erato*) was enriched in binding sites for *Mothers against dpp* (*Mad*; FBgn0011648), *buttonhead* (*btd*; FBgn0000233), *schnurri* (*shn*; FBgn0003396), and *48 related 1* (*Fer1*; FBgn0037475). TFs with known functions in neuron development are indicated with an asterisk. P-value is the significance of enrichment as obtained by the MEME analysis **(B.)** Homology of identified enriched motifs to *D. melanogaster* motifs of TFs with known functions in neuron development.

Finally, using the ATAC-seq data of the wing tissue as a control, we looked for head-specific chromatin changes within lineage-specific TE insertions. The tissue-specific accessibility of these TE insertions would provide strong indications that these SVs interact with tissue-specific factors and could provide strong candidates as targets of adaptive evolution. We identified 24 head-specific ATAC-seq peaks within a lineage-specific TE insertion that were not accessible in wing tissues (Figure 6; Table S3). Of these, 2, 4 and 18 were specific to *H. charithonia, H. erato* and *H. melpomene*, respectively. Five were located less than 50 kb from genes with known functions in nervous system development in *Drosophila*. Interestingly, in *H. melpomene*, this included the gene *sloppy paired 2* (*slp2*) that also showed TF binding site enrichment in lineage-specific ATAC-seq peaks within a TE.

**Figure 6.**
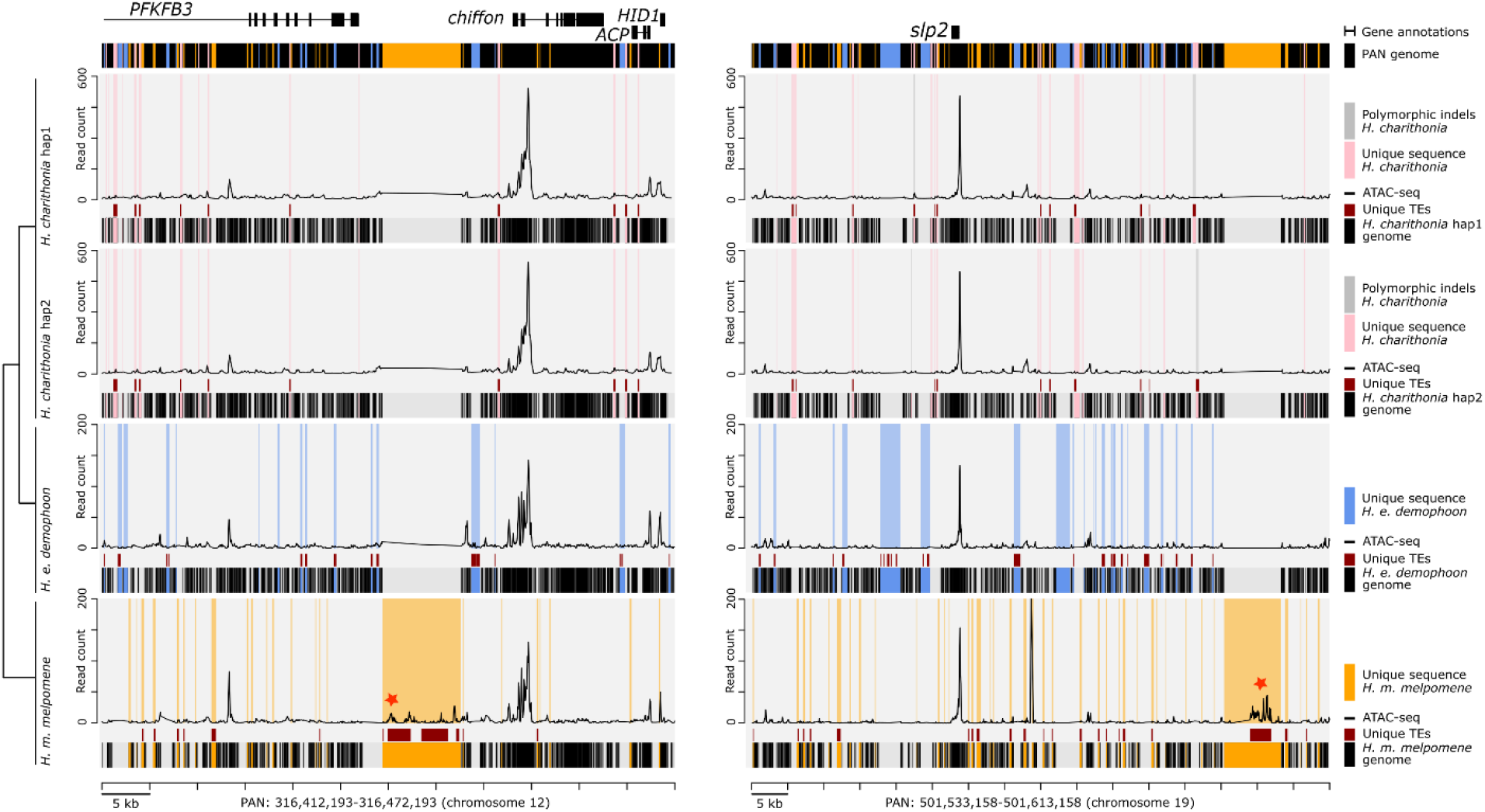
Example intervals of the pan-genome assembly of *H. charithonia* (pink), *H. erato* (blue) and *H. melpomene* (orange) with alignment of lineage-specific genome sequences, transposable element (TE) annotations and ATAC-seq profiles in the pan-genome coordinate space. The plots show an illustrative interval of the pan-genome assembly near the gene *chiffon* and *slp2* that highlights sequences present in each of the genomes relative to the pan-genome (black shading underneath each of the graphs), lineage-specific sequences in each of the genomes (pink, blue and orange shading in graphs), TEs that overlap with lineage-specific sequences (dark red), and ATAC-seq profiles for head tissue (average of two biological replicates). Gray shading in the *H. charithonia* haplotype 2 (hap2) graph indicates an indel in the genome of a single *H. charithonia* individual. Red stars indicate ATAC-seq peaks with head-specific accessibility (compared to wing tissue) that intersect with a lineage-specific TE insertion.

## Conclusion

In *Heliconius*, the extent of SV within and between species has been previously limited to studies of the repetitive sequence content within individual reference genomes (Lavoie et al. 2013; Ray et al. 2019), co-linearity of genomes (Davey et al. 2017; Cicconardi et al. 2021), structural rearrangements in a ‘supergene’ related to a color pattern polymorphism (Joron et al. 2011; Edelman et al. 2019), and of duplications that likely underestimated the extent of SV due to stringent confidence cutoffs needed when using short-read sequences (Pinharanda et al. 2017). Our approach combined three high-quality *Heliconius* genome assemblies with a pan-genome alignment to quantify the extensive uniqueness between these genomes due to SVs. For example, genome wide nucleotide diversity (π) obtained from SNPs was 0.0098 within *H. charithonia* and *D*_*XY*_ (average pairwise nucleotide differences) ranged from 0.10 between *H. charithonia* and *H. erato* to 0.15 between *H. charithonia* and *H. melpomene*. This suggests an average sequence divergence of 0.98% between the haplotypes of *H. charithonia* and 10 to 15% of sequence divergence between homologous parts of the genomes of these species. In contrast, SV analysis demonstrated that an additional 18.0% of the genome of *H. charithonia* included indel sequences and up to 43.5% and 62.7% of additional genomic differences between *H. charithonia* and *H. erato* and *H. melpomene*, respectively, resulted from SVs. Next, using ATAC-seq data, we assessed the extent to which differences in chromatin accessibility resulted from SVs and TE insertions. We observed that out of the 515,884 SVs identified as lineage-specific TE insertions, only 0.56% were associated with changes in chromatin accessibility between species. However, out of the 31,066 identified lineage-specific changes in chromatin accessibility, 30.4% were within SVs and 9.4% were characterized as lineage-specific TEs. While we did not find any indication of recent adaptive evolution through selective sweep analysis, our observations indicate an important potential of TEs in generating genetic variation with functional effects through changes in chromatin state and potentially the regulation of nearby genes. In support with this, lineage-specific TEs that underlie changes in chromatin accessibility included 21 enriched motifs homologous to *Drosophila* TF binding sites and 24 had head-specific accessibility compared to wing tissues and provide strong candidates as targets of adaptive evolution.

Recent examples that characterized SVs and, in particular, TE insertions as the mutational changes underlying adaptive phenotypic changes are accumulating. For example, in the bird genus *Corvus*, adaptive evolution of plumage patterning, a pre-mating isolation trait, was found to be the result of a TE insertion that reduced the expression of the *NDP* gene (Weissensteiner et al. 2020). Together with examples of adaptive phenotype changes associated with TEs in Lepidoptera (Livraghi et al. 2021; Van’t Hof 2016), our pan-genome strategy that allowed to efficiently study the genomic distribution of SVs and the evolutionary factors that affect them, suggest that TE insertions coupled to gene regulation may be an underappreciated source of variation for natural selection to act upon. We expect that the accumulation of high-quality genome assemblies thanks to long-read sequencing technologies will continue to improve the identification of SVs and highlight their importance in generating adaptive genetic variation.

## Materials and methods

### Heliconius charithonia haploid genome assemblies

For *Heliconius charithonia*, we extracted high-molecular-weight DNA from a flash frozen pupa obtained from a wild-caught female sampled in San Juan, Puerto Rico using QIAGEN Inc. Genomic-tip 100/G. Library preparation using 10X Chromium technology for linked reads (10x Genomics, San Francisco, USA) and Illumina sequencing was carried out by Novogene Co., Ltd, which generated 44.9 Gb for a target coverage of 100X. We assembled the linked-read sequencing data using the Supernova 2.1.1 assembler (Weisenfeld et al. 2014) using the default recommended settings and a maximum number of reads of 200 million. Raw assembly outputs were transformed to fasta format using the pseudohap2 option to generate two parallel pseudo-haplotypes from the diploid genome. Quality control of the *H. charithonia* genome was performed using genome-wide statistics calculated on the phase blocks, synteny with the *H. melpomene* v2.5 genome using Tigmint v1.2.3 (Jackman et al. 2018), and using BUSCO to assess genome assembly and annotation completeness (Simão et al. 2015). Fragmented *H. charithonia* scaffolds were ordered with Tigmint using synteny with the *H. melpomene* v2.5 genome.

### Pan-genome alignment

In comparison to using a single genome as a reference, a pan-genome represents a composite of different genomes and serves as a global reference with which to make comparisons between genomes (e.g., conservation and unique sequences) or genome features (e.g., gene and TE annotations). We aligned the two newly assembled haploid *H. charithonia* genomes with the *H. e. demophoon* and *H. m. melpomene* genome using seq-seq-pan (Jandrasits et al. 2018). Seq-seq-pan extends the functionality of the multiple genome aligner progressiveMauve (Darling et al. 2010) by constructing a composite consensus or pan-genome that includes both homologous sequences or locally collinear blocks (LCBs) as well as lineage-specific (non-homologous) sequences in each of the genomes. This pan-genome is then used as the reference coordinates space for the multi genome alignment which can then include sequences specific to any of the genomes. We used the *H. e. demophoon* v1 reference genome as the first genome in the genome list so that the resulting pan-genome alignment would be ordered according to the *H. e. demophoon* reference. This resulted in a pan-genome sequence with a total length of 659,350,588 bp. To avoid spurious feature mappings (i.e., TEs and ATAC-seq peaks), we removed scaffolds that have not been linked to chromosome positions in *H. e. demophoon* by cutting the pan-genome alignment at the end of chromosome 21 (position 578,665,626 in the alignment). The absence and presence of genome sequences in each of the genomes relative to the pan-genomes was assessed with a custom Python script generating a bed file of start and end positions of presence and absence data. These bed files were used to identify lineage-specific or homologous sequences between genomes using bedtools v2.27.1 (Quinlan and Hall 2010). Lineage-specific sequences were obtained by first recording sequence coordinates of each genome relative to the pan-genome using a custom Python script and intersecting these coordinates of each genome against a merged library of sequence coordinates of all other genomes using bedtools.

### Transposable Element (TE) annotation and analysis

To identify TEs, we used a two-stage strategy combining the programs RepeatModeler2 (Flynn et al. 2020) and RepeatMasker (Tarailo-Graovac and Chen 2009) using available curated TE libraries as well as novel TE discovery. In the first stage, RepeatModeler 2.0.1 was run on the four genomes for *de novo* identification of TEs, to classify them into families, and merge the results into a single library. We used the Perl script “cleanup_nested.pl” from the LTR_retriever package (Ou and Jiang 2018) with default parameters to reduce redundant and nested TEs. The TE library was then filtered to eliminate all sequences shorter than 200 bp and all sequences that matched any non TE-related genes using Blast2GO (Conesa et al. 2005) with the insect-only default library (non-redundant protein sequence nr v5). Finally, the filtered TEs were matched with the *Heliconius* specific TE library from Ray *et al*. (2019) using Blast2GO. This library was produced with *de novo* TE annotations of 19 Heliconiinae and including *H. erato* and *H. melpomene*. The remaining sequences with a TE annotation from RepeatModeler that did not match the *Heliconius* specific TE library from Ray *et al*. (2019) were analyzed with different strategies appropriate for the transposon type. First, the putative autonomous elements (DNA, LTR, and LINE) were analyzed with Blast2GO against the insect-only default library. DNA and LTR elements had to have at least a TE-derived transposase and/or match with other DNA/LTR elements. The LINE required the presence of a reverse transcriptase. Second, the putative SINEs were searched in SINEbase (Vassetzky and Kramerov 2013) and accepted only if at least one of their parts (head, body, tail) matched with a SINE element in the database. Third, the putative Helitrons were identified using DeepTE with the parameters -sp M -m M -fam ClassII (Yan et al. 2020). TEs identified as Helitrons were then scanned with CENSOR (Kohany et al. 2006) to confirm their origin. From these analyses we annotated an additional 93 TEs compared to Ray *et al*. (2019). These TEs were labelled as “putative TEs” and were added to the library from Ray *et al*. (2019) to obtain the final library.

In the second stage, we used the non-redundant library as a custom library in RepeatMasker 4.1.0 to annotate the Tes within our genomes. The RepeatMasker results were cleaned with “one code to find them all” (Bailly-Bechet et al. 2014). This script combines fragmented RepeatMasker hits into complete TE copies and solves ambiguous cases of nested TE. We identified TE families that have been differentially active between phylogenetic branches using a chi-square test with false discovery rate correction. We characterized temporal variation of Metulj-7 and Metulj-m51, two TEs that showed the strongest temporal changes in activity, using the percent of divergence compared to the TE library reference sequence obtained from RepeatMasker, corrected with the Jukes-Cantor model. Finally, TE fragmentation was calculated based on the total length of each element recovered from the reference library.

### ATAC-seq library preparation

ATAC-seq libraries were constructed as in Lewis and Reed (2019), a protocol modified from Buenrostro et al. (2013), with minor modifications. *H. melpomene rosina* and *H. erato demophoon* butterflies were collected in Gamboa, Panama; *H. charithonia* butterflies were collected in San Juan, Puerto Rico. Two caterpillars of each species were reared on their respective host plants and allowed to grow until the wandering stage at 5th instar. Live larvae were placed on ice for 1-2 minutes and then pinned and dissected in 1X ice cold PBS. Using dissection scissors, the head was removed, and incisions were performed between the mandibles and at the base of the vertexes. Fine forceps were then used to remove the head cuticle to expose the tissue below. The brain and eye-antennal tissue was subsequently dissected out, by removing the remaining cuticle still attached to the tissue. Similarly, developing wings were dissected from the 5th instar caterpillars and the left and right forewing and left and right hindwing were pooled, respectively.

The tissues were then submerged in 350 μl of sucrose solution (250mM D-sucrose, 10mM Tris-HCl, 1mM. MgCl2, 1x protease inhibitors) inside 2 ml dounce homogenizers for tissue homogenization and nuclear extraction. After homogenizing the tissue on ice, the resulting cloudy solution was centrifuged at 1000 rcf for 7 minutes at 4 °C. The pellet was then resuspended in 150 μl of cold lysis buffer (10mM Tris-HCl, 10mM NaCl, 3mM MgCl2, 0.1% IGEPAL CA-630 (SigmaAldrich), 1x protease inhibitors) to burst the cell membranes and release nuclei into the solution. Samples were then checked under a microscope with a counting chamber following each nuclear extraction, to confirm nuclei dissociation and state and to assess the concentration of nuclei in the sample. Finally based on these observations a calculation to assess the number of nuclei, and therefore DNA, to be exposed to the transposase was performed. This number was fixed on 400,000, as it is the number of nuclei required to obtain the same amount of DNA from a ∼0.4 Gb genome, such as that of *H. erato* and *H. charithonia*, as is contained in 50,000 human nuclei – the amount of DNA for which ATAC-seq is optimized (Buenrostro et al. 2013).

For *H. melpomene* this number was 500,333, where the genome size of *H. melpomene* is 0.275 Gb. For this quality control, a 15 μl aliquot of nuclear suspension was stained with trypan blue, placed on a hemocytometer and imaged at 64x. After confirmation of adequate nuclear quality and assessment of nuclear concentration, a subsample of the volume corresponding to 400,000 nuclei (*H. erato* and *H. charithonia*) and 500,333 (*H. melpomene*) was aliquoted, pelleted 1000 rcf for 7 minutes at 4 °C and immediately resuspended in a transposition mix, containing Tn5 enzyme (Illumina DNA Prep) in a transposition buffer. The transposition reaction was incubated at 37 °C for exactly 30 minutes. A PCR Minelute Purification Kit (Qiagen) was used to interrupt the tagmentation and purify the resulting tagged fragments, which were amplified using custom-made Nextera primers and a NEBNext High-fidelity 2x PCR Master Mix (New England Labs). The amplified libraries were sequenced as 37 to 76 bp paired-end fragments with NextSeq 500 Illumina technology at the Sequencing and Genomics Facility of the University of Puerto Rico (Table S4).

### ATAC-seq data analysis

Raw Illumina reads were filtered for adapters and quality using Trimmomatic v0.39 (Bolger et al. 2014). Filtered reads for each sample were then mapped to their respective reference genome using bowtie2 v2.2.6 (Langmead and Salzberg 2013) using default parameters. We used; samtools v1.2 (Li et al. 2009) to sort mapped reads and only retain reads with a mapping phred score higher than 20 (-q 20) and that were uniquely mapped and properly oriented (-f 0×02). PCR duplicates were identified and removed using picard-tools v2.5 (http://picard.sourceforge.net).

ATAC-seq peak intervals were called on the mapped reads (bam files) of each sample using the MACS2 ‘callpeak’ command with –g set to the respective reference genome size and –shift set to -100 and – extsize set to 200 (Zhang et al. 2008). Peaks were only retained if they occurred in both replicates with a reciprocal minimal 25% overlap, as determined with bedtools intersect function. The function ‘multicov’ from bedtools was used to obtain read counts within ATAC-seq peaks. These read counts were used to obtain library size scaling factors using the function ‘estimateSizeFactors’ from the R package DESeq2 (Love et al. 2014). Next, bam files were converted to bedgraphs using the bedtools function ‘genomecov’ and scaled using the size scaling factors. Mean ATAC-seq traces for each species were obtained from the two replicate samples using wiggletools (Zerbino et al. 2014). Differential accessibility between head and wing tissues was tested in each species using DESeq2 (Love et al. 2014) with an adjusted p-value smaller than 0.05 and fold change larger than 1.

### Feature mapping to pan-genome coordinates and comparisons

Features, including genome sequences that are lineage-specific, TE annotations from RepeatMasker, gene annotations (obtained from *H. e. demophoon*), and ATAC-seq peaks from MACS2 were compared after converting their genome coordinates to pan-genome coordinates. This was done by first using the ‘map’ utility of the seq-seq-pan software (Jandrasits et al. 2018) and custom scripts. Features that overlapped with scaffold starts or ends in any of the genomes were masked using bedtools ‘subtract’ (-A) to avoid including results from fragmented or missing sequences. Next, lineage-specific sequences were intersected with TE annotations and ATAC-seq peaks using bedtools ‘intersect’ with a minimum overlap of 25% (-f 0.25). Lineage-specific sequences in one of the genomes that did not match a TE annotation were identified as duplications when identifying a blast hit with a similarity higher than 70% elsewhere in the genome using blast v2.10.0.

### Feature distribution

We measured the genomic distance along the pan-genome of lineage-specific sequences, TEs and ATAC-seq peaks from the closest transcription start site (TSS) of a gene using the function ‘annotatePeaks’ from the software suite Homer (Heinz et al. 2010). Each distribution was compared with that of 100,000 random positions with a pairwise Wilcoxon test. For each distribution pair an overlapping index was measured, using the R package *overlapping* v1.6 (Pastore 2018).

### Motif enrichment

Differential motif enrichment analysis was performed for ATAC-seq peaks that overlapped with lineage-specific TEs using the STREME tool from the MEME suite (Machanick & Bailey 2011; Bailey 2020). This was done for four phylogenetic comparisons: *H. charithonia* compared to *H. erato, H. charithonia* compared to *H. melpomene, H. erato* compared to *H. melpomene*, and *H. melpomene* compared to *H. erato*. As a background model, we constructed a custom dataset including a combined set of lineage-specific TEs without ATAC-seq peaks from the phylogenetic comparisons. Motifs with a p-value smaller than 0.001 were analyzed with Tomtom from the MEME-suite to identify motifs similar transcription factor binding sites in *Drosophila melanogaster* (Gupta et al. 2007).

### Historical population demography

Changes in historical population sizes from individual genome sequences were inferred using the pairwise sequentially Markovian coalescent (PSMC) as implemented in MSMC (Schiffels and Durbin 2014). Genotypes were inferred using SAMTOOLS v0.1.19 (Li et al. 2009) from reads mapped to the respective reference genomes using BWA v0.7 (Li and Durbin 2010). This involved a minimum mapping (-q) and base (-Q) quality of 20 and adjustment of mapping quality (-C) 50. A mask file was generated for regions of the genome with a minimum coverage depth of 30 and was provided together with heterozygosity calls to the MSMC tool. MSMC was run on heterozygosity calls from all contiguous scaffolds longer than 500 kb, excluding scaffolds on the Z chromosome. We scaled the PSMC estimates using a generation time of 0.25 years and a mutation rate of 2e-9 as estimated for *H. melpomene* (i.e., spontaneous *Heliconius* mutation rate corrected for selective constraint (Keightley et al. 2014; Martin et al. 2015)). We obtained whole-genome resequencing reads for *H. e. demophoon* and *H. m. melpomene* from two individuals each from Panama (SAMN05224182, SAMN05224183, SAMEA1919255, and SAMEA1919258 from Van Belleghem et al. 2018). For *H. charithonia*, we obtained resequencing data for one sample from Panama (SAMN05224120 from Van Belleghem et al. 2017) and two samples from Puerto Rico (SAMN05224121 from Van Belleghem *et al*. (2017) and one using the 10X linked-read sequencing data used for the genome assembly from the Puerto Rican population).

### Signatures of selective sweeps

SweepFinder2 (Degiorgio et al. 2016) was used to detect signatures of selective sweeps in genomic regions with ATAC-seq peaks with lineage-specific TEs. Genotypes from 10 *H. erato demophoon* and 10 *H. melpomene rosina* individuals from Panamanian populations were obtained from Van Belleghem et al. (2018). Allele counts for biallelic SNPs were generated using a custom Python script. SNPs were polarized using *H. hermathena* and *H. numata* for the *H. erato* and *H. melpomene* population, respectively. SweepFinder2 was run using default settings and set to test SNPs every 2000 bp (-sg 2000).

## Data accessibility

This Whole Genome Shotgun project for the *H. charithonia* pseudo-haplotypes has been deposited at DDBJ/ENA/GenBank under the accession JAKFBP000000000. The version described in this paper is version JAKFBP010000000. The raw 10X chromium sequencing reads for the genome assembly are deposited on SRA (SAMN24661992). ATAC-seq raw reads are available under NCBI Bioproject ID PRJNA795145 (SAMN24689923-SAMN24689940). Code for analyses is available at https://github.com/StevenVB12/Genomics.

## Acknowledgements

We thank Christine Jandrasits for advice in using seq-seq-pan, Markus Möst for help with running SweepFinder2 and Simon H. Martin for help in interpreting chromosomal indel diversity patterns. We also thank Yadira Ortiz-Ruiz and Silvia Planas from the Sequencing and Genomics Facility of the University of Puerto Rico-Rio Piedras for their assistance with DNA extractions and ATAC-seq library preparation and sequencing. This work was supported by a National Institutes of Health-4 NIGMS COBRE Phase 2 Award – Center for Neuroplasticity at the University of Puerto Rico (Grant No. 1P20GM103642) to SMVB, a Puerto Rico Science, Technology & Research Trust catalyzer award (#2020-00142) to SMVB and RP, and an NSF EPSCoR RII Track-2 FEC (grant no. OIA 1736026), an NSF IOS 1656389, and a Fondo Institucional para la Investigacio’n (FIPI), Universidad de Puerto Rico - Recinto de Río Piedras, Decanato de Estudios Graduados e Investigación to RP. For sequencing and computational resources, we thank the University of Puerto Rico, the Puerto Rico INBRE Grant P20 GM103475 from the National Institute for General Medical Sciences (NIGMS), a component of the National Institutes of Health (NIH), and the Bioinformatics Research Core of the INBRE. Its contents are solely the responsibility of the authors and do not necessarily represent the official view of NIGMS or NIH.

## SUPPLEMENTAL MATERIALS

**Table S1.** Genome assembly sizes, N50 and BUSCO completeness. Busco: C = complete, S = complete and single-copy, D = complete and duplicated, F = fragmented, M = missing.

| Species | Assembly size (Mb) | N50 (kb) | BUSCO completeness |
| --- | --- | --- | --- |
| *H. melpomene* | 275.3 | 14,309 | C:99.3% [S:98.3%, D:1.0%], F:0.3%, M:0.4% |
| *H. erato* | 382.8 | 10,689 | C:98.8% [S:97.6%, D:1.2%], F:0.2%, M:1.0% |
| *H. charithonia* (hap 1) | 355.2 | 11,106 | C:98.9% [S:97.9%, D:1.0%], F:0.8%, M:0.4% |
| *H. charithonia* (hap 2) | 361.5 | 11,118 | C:98.2% [S:97.4%, D:0.8%], F:0.8%, M:1.0% |
| Homologous (pan-genome) | 138.6 | NA | C:94.9% [S:94.8%, D:0.1%], F:1.4%, M:3.7% |

**Table S2.** Lineage-specific sequence composition. Lineage-specific (unique) sequences were characterized as being 1 bp, shorter than 20 bp (< 20 bp), larger than 1,000 bp, Transposable Elements (TE), or duplicated. % genome = percentage of total length of genome, % count = percentage of unique sequence count, % unique = percentage of unique sequence base pairs.

|  | Unique<br>sequence in Mb<br>(% genome) | 1 bp in Mb<br>(% genome /% count<br>/% unique) | < 20 bp in Mb<br>(% genome /% count<br>/% unique) | > 1000 bp in Mb<br>(% genome /% count<br>/% unique) | TE<br>(%<br>unique) | Dupl.<br>(%<br>unique) | Total<br>identified<br>(% unique) |
| --- | --- | --- | --- | --- | --- | --- | --- |
| *H. charithonia*<br>(indels) | 72.7<br>(18.0) | 0.74<br>(0.18 / 25.78 / 1.92) | 8.04<br>(2.00 / 95.49 / 20.88) | 20.46<br>(5.07 / 0.13 / 53.09) | 42.36 | 5.21 | 63.79 |
| *H. charithonia*<br>(compared to *H. erato*) | 175.2<br>(43.45) | 0.16<br>(0.04 / 5.76 / 1.57) | 11.02<br>(2.73 / 81.96 / 10.64) | 33.74<br>(8.37 / 0.35 / 32.58) | 52.22 | 6.17 | 69.78 |
| *H. erato*<br>(compared to *H. charithonia*) | 151.21<br>(39.49) | 0.21<br>(0.05 / 7.11 / 0.14) | 11.44<br>(2.99 / 77.91 / 67.57) | 63.98<br>(16.71 / 0.71 / 42.31) | 52.85 | 10.66 | 70.03 |
| *H. charithonia*<br>(compared to *H. melpomene*) | 252.7<br>(62.67) | 0.27<br>(0.07 / 6.90 / 0.15) | 12.83<br>(3.18 / 67.50 / 7.09) | 34.82<br>(8.64 / 0.27 / 19.24) | 42.50 | 9.10 | 58.69 |
| *H. erato*<br>(compared to *H. melpomene*) | 222.12<br>(58.02) | 0.17<br>(0.04 / 4.48 / 0.08) | 12.93<br>(3.38 / 64.67 / 5.82) | 65.39<br>(17.08 / 0.56 / 29.44) | 49.48 | 7.12 | 61.63 |
| *H. melpomene*<br>(compared to *H. charithonia*) | 113.5<br>(41.24) | 0.17<br>(0.06 / 6.14 / 0.15) | 10.27<br>(3.73 / 75.87 / 9.06) | 29.18<br>(10.60 / 0.44 / 25.73) | 47.34 | 5.29 | 61.83 |
| *H. melpomene*<br>(compared to *H. erato*) | 106.83<br>(38.82) | 0.13<br>(0.05 / 4.93 / 0.12) | 9.64<br>(3.50 / 78.34 / 9.02) | 29.01<br>(10.54 / 0.46 / 27.15) | 47.34 | 5.29 | 61.83 |

**Table S3.**
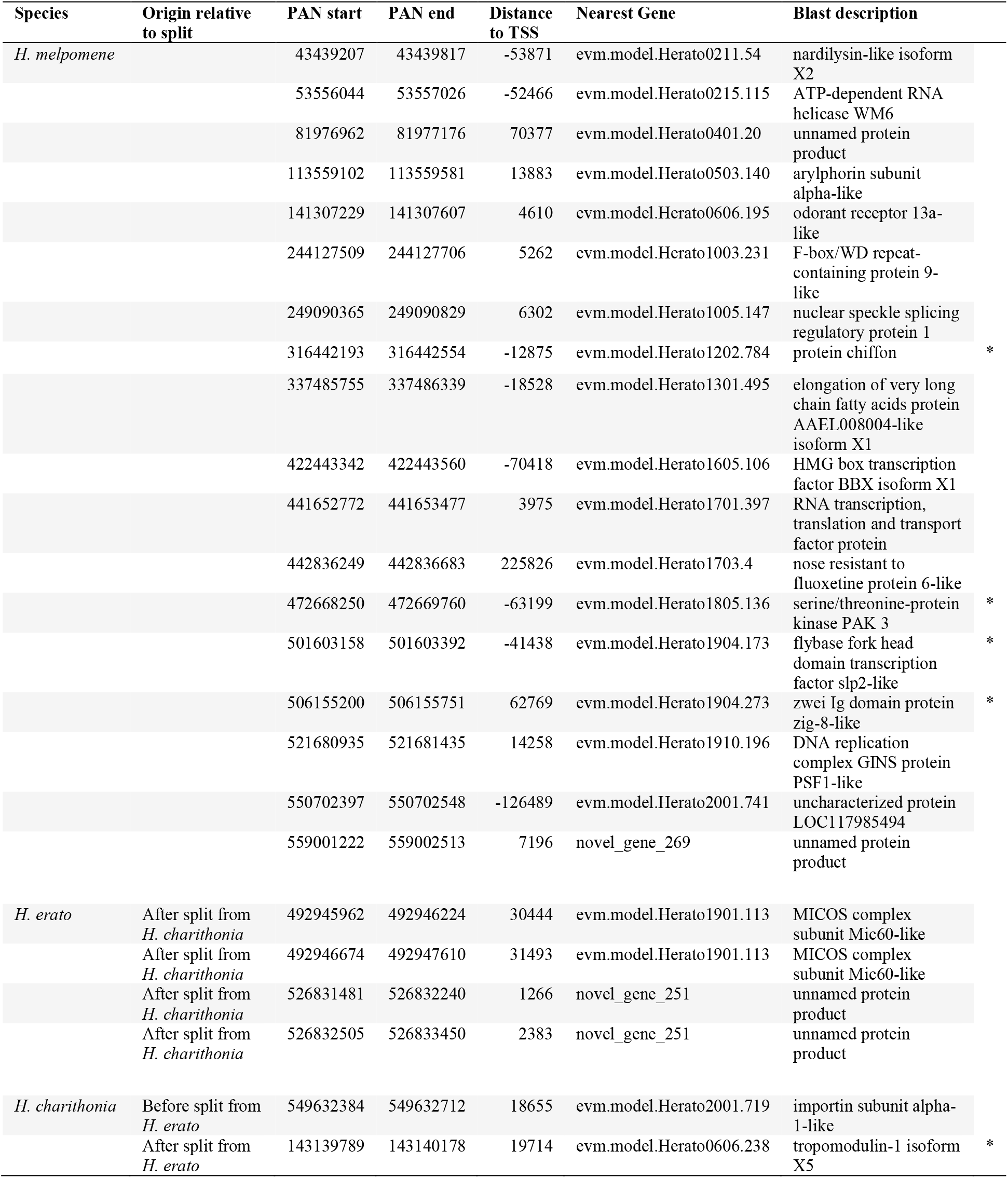
Genes near lineage-specific ATAC-seq peaks with head-specific accessibility and that associate with lineage-specific TE insertions. Rows indicated with an asterisk are peaks near genes with a known function related to eye or neuron development. Note that origin relative to their split can only be presented for *H. erato* and *H. charithonia*.

| Species | Origin relative to split | PAN start | PAN end | Distance to TSS | Nearest Gene | Blast description |  |
| --- | --- | --- | --- | --- | --- | --- | --- |
| *H. melpomene* |  | 43439207 | 43439817 | -53871 | evm.model.Herato0211.54 | nardilysin-like isoform X2 |  |
|  |  | 53556044 | 53557026 | -52466 | evm.model.Herato0215.115 | ATP-dependent RNA helicase WM6 |  |
|  |  | 81976962 | 81977176 | 70377 | evm.model.Herato0401.20 | unnamed protein product |  |
|  |  | 113559102 | 113559581 | 13883 | evm.model.Herato0503.140 | arylphorin subunit alpha-like |  |
|  |  | 141307229 | 141307607 | 4610 | evm.model.Herato0606.195 | odorant receptor 13a-like |  |
|  |  | 244127509 | 244127706 | 5262 | evm.model.Herato1003.231 | F-box/WD repeat-containing protein 9-like |  |
|  |  | 249090365 | 249090829 | 6302 | evm.model.Herato1005.147 | nuclear speckle splicing regulatory protein 1 |  |
|  |  | 316442193 | 316442554 | -12875 | evm.model.Herato1202.784 | protein chiffon | * |
|  |  | 337485755 | 337486339 | -18528 | evm.model.Herato1301.495 | elongation of very long chain fatty acids protein AAEL008004-like isoform X1 |  |
|  |  | 422443342 | 422443560 | -70418 | evm.model.Herato1605.106 | HMG box transcription factor BBX isoform X1 |  |
|  |  | 441652772 | 441653477 | 3975 | evm.model.Herato1701.397 | RNA transcription, translation and transport factor protein |  |
|  |  | 442836249 | 442836683 | 225826 | evm.model.Herato1703.4 | nose resistant to fluoxetine protein 6-like |  |
|  |  | 472668250 | 472669760 | -63199 | evm.model.Herato1805.136 | serine/threonine-protein kinase PAK 3 | * |
|  |  | 501603158 | 501603392 | -41438 | evm.model.Herato1904.173 | flybase fork head domain transcription factor slp2-like | * |
|  |  | 506155200 | 506155751 | 62769 | evm.model.Herato1904.273 | zwei Ig domain protein zig-8-like | * |
|  |  | 521680935 | 521681435 | 14258 | evm.model.Herato1910.196 | DNA replication complex GINS protein PSF1-like |  |
|  |  | 550702397 | 550702548 | -126489 | evm.model.Herato2001.741 | uncharacterized protein LOC117985494 |  |
|  |  | 559001222 | 559002513 | 7196 | novel_gene_269 | unnamed protein product |  |
| *H. erato* | After split from *H. charithonia* | 492945962 | 492946224 | 30444 | evm.model.Herato1901.113 | MICOS complex subunit Mic60-like |  |
|  | After split from *H. charithonia* | 492946674 | 492947610 | 31493 | evm.model.Herato1901.113 | MICOS complex subunit Mic60-like |  |
|  | After split from *H. charithonia* | 526831481 | 526832240 | 1266 | novel_gene_251 | unnamed protein product |  |
|  | After split from *H. charithonia* | 526832505 | 526833450 | 2383 | novel_gene_251 | unnamed protein product |  |
| *H. charithonia* | Before split from *H. erato* | 549632384 | 549632712 | 18655 | evm.model.Herato2001.719 | importin subunit alpha-1-like |  |
|  | After split from *H. erato* | 143139789 | 143140178 | 19714 | evm.model.Herato0606.238 | tropomodulin-1 isoform X5 | * |

**Table S4.** ATAC-seq samples. Number of reads are quality filtered and properly paired.

| Sample ID | Tissue | Species | Locality | Accession # | # reads |
| --- | --- | --- | --- | --- | --- |
| CH_head2 | Head | *H. charithonia* | Puerto Rico | SAMN24689923 | 14,045,582 |
| CH_head6 | Head | *H. charithonia* | Puerto Rico | SAMN24689924 | 36,694,766 |
| BR12 | Head | *H. erato demophoon* | Panama | SAMN24689925 | 18,019,744 |
| BR14 | Head | *H. erato demophoon* | Panama | SAMN24689926 | 57,988,466 |
| BR11 | Head | *H. melpomene rosina* | Panama | SAMN24689927 | 10,214,746 |
| M4_head | Head | *H. melpomene rosina* | Panama | SAMN24689928 | 45,921,776 |
| CH_FW2 | Forewing | *H. charithonia* | Puerto Rico | SAMN24689929 | 8,605,992 |
| CH_FW7 | Forewing | *H. charithonia* | Puerto Rico | SAMN24689930 | 23,397,866 |
| LI7_demophoon_FW | Forewing | *H. erato demophoon* | Panama | SAMN24689931 | 17,294,536 |
| FW-pboy | Forewing | *H. erato demophoon* | Panama | SAMN24689932 | 49,257,240 |
| LI14_rosina_FW | Forewing | *H. melpomene rosina* | Panama | SAMN24689933 | 23,449,240 |
| M4_FW | Forewing | *H. melpomene rosina* | Panama | SAMN24689934 | 24,353,808 |
| CH_HW2 | Hindwing | *H. charithonia* | Puerto Rico | SAMN24689935 | 26,910,064 |
| CH_HW7 | Hindwing | *H. charithonia* | Puerto Rico | SAMN24689936 | 21,029,072 |
| LI7_demophoon_HW | Hindwing | *H. erato demophoon* | Panama | SAMN24689937 | 14,631,856 |
| LB_42 | Hindwing | *H. erato demophoon* | Panama | SAMN24689938 | 26,635,566 |
| LI14_rosina_HW | Hindwing | *H. melpomene rosina* | Panama | SAMN24689939 | 23,176,622 |
| M4_HW | Hindwing | *H. melpomene rosina* | Panama | SAMN24689940 | 36,959,108 |

**Figure S1.**
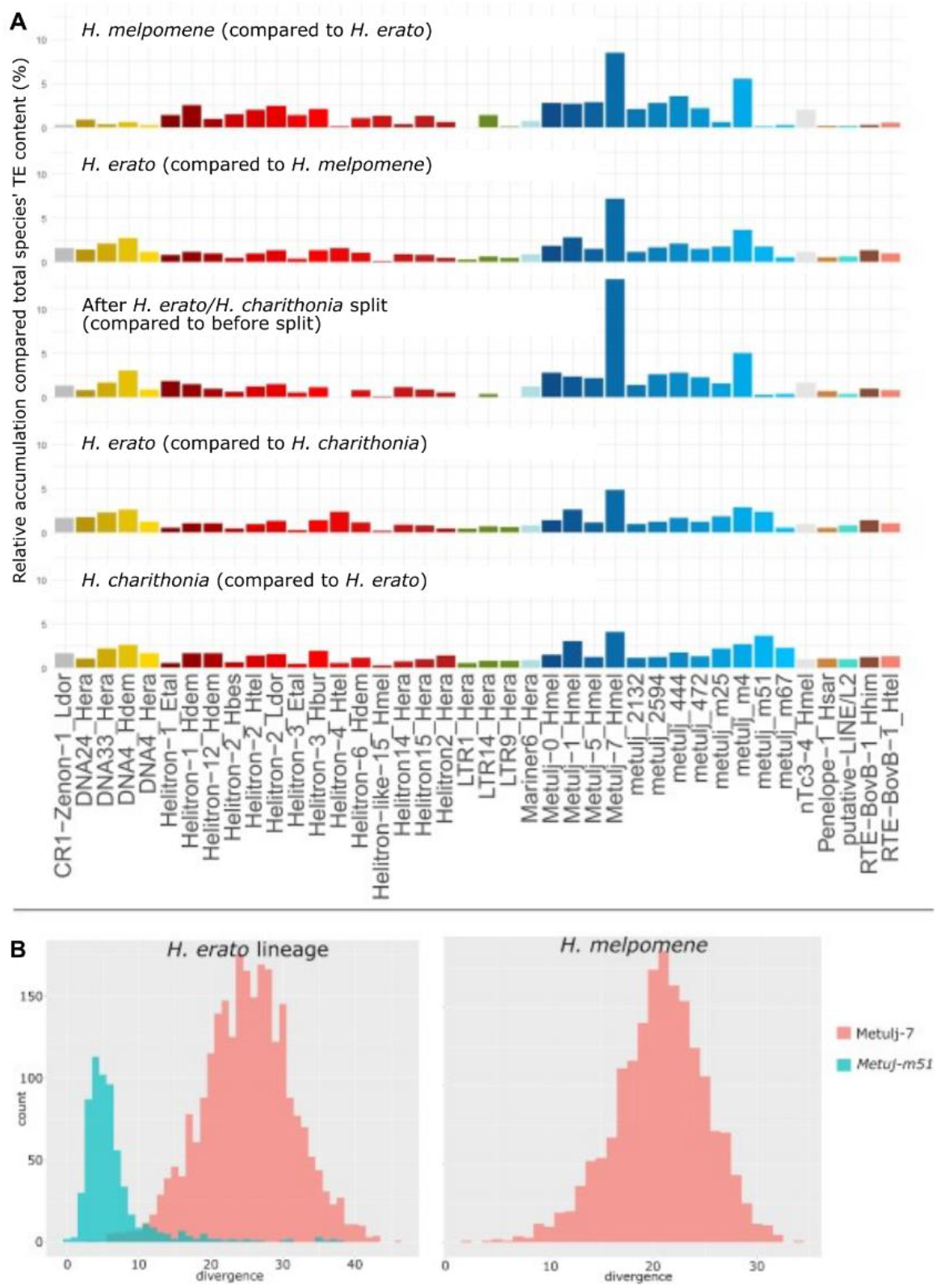
Lineage specific transposable element activity. **(A.)** Sccumulation of the 40 most active TE families in each lineage (values are percentages among the total lineage-specific TEs in the species). These families were obtained by grouping the 20 most active families for each lineage comparison (see Figure 3). (**B.)** Divergence distribution plots of Metulj-7 and Metulj-m51 represent times of activity in *H. erato* and *H. melpomene*. The x-axis represents the divergence (percentage) from the reference sequence corrected with the Jukes-Cantor model, the y-axis represents the sequence count.

**Figure S2.**
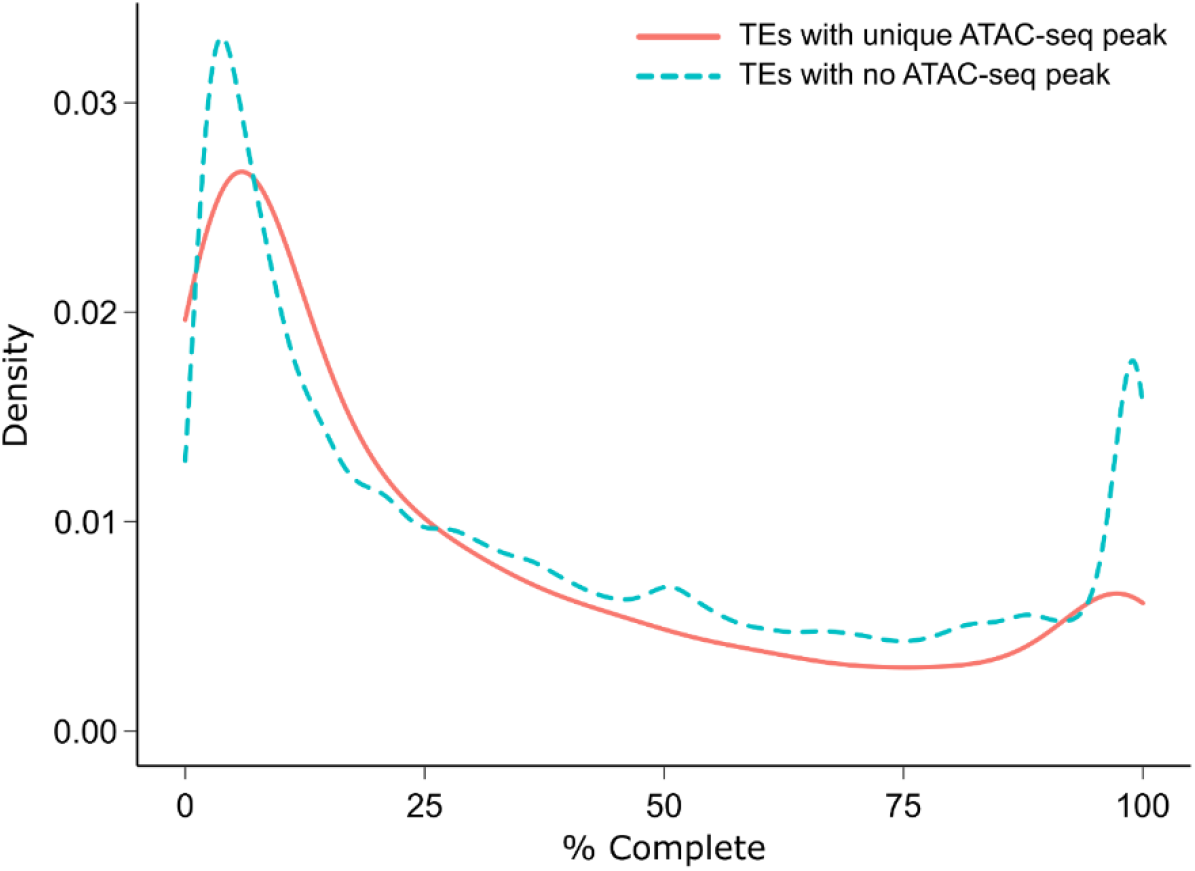
Completeness of Transposable Elements (TEs) for TEs associated with lineage-specific ATAC-peaks and TEs with no ATAC-seq peak association. TE completeness was calculated based on the total length of each element recovered from the reference library.

**Figure S3.**
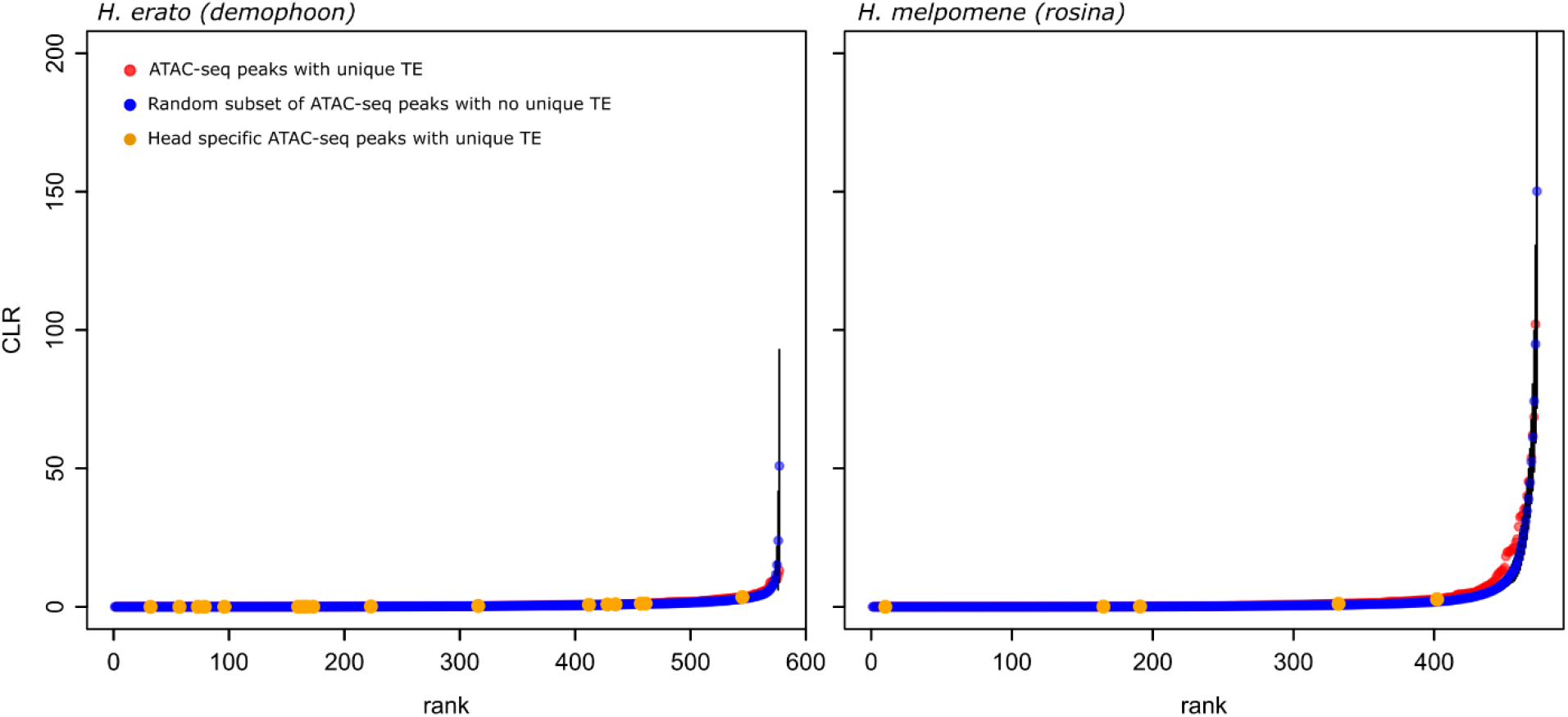
Ranked SweepFinder2 Composite Likelihood Ratio (CLR) values for ATAC-seq peaks with and without lineage-specific Transposable Element (TE). ATAC-seq peaks with lineage-specific TE (red) do not show greater signatures of selective sweeps than expected from random subsets of ATAC-seq peaks with no lineage-specific TE (blue) in both populations of *H. erato demophoon* (left) and *H. melpomene rosina* (right). Head-specific ATAC-seq peaks with a lineage-specific TE are indicated in orange.

**Figure S4.**
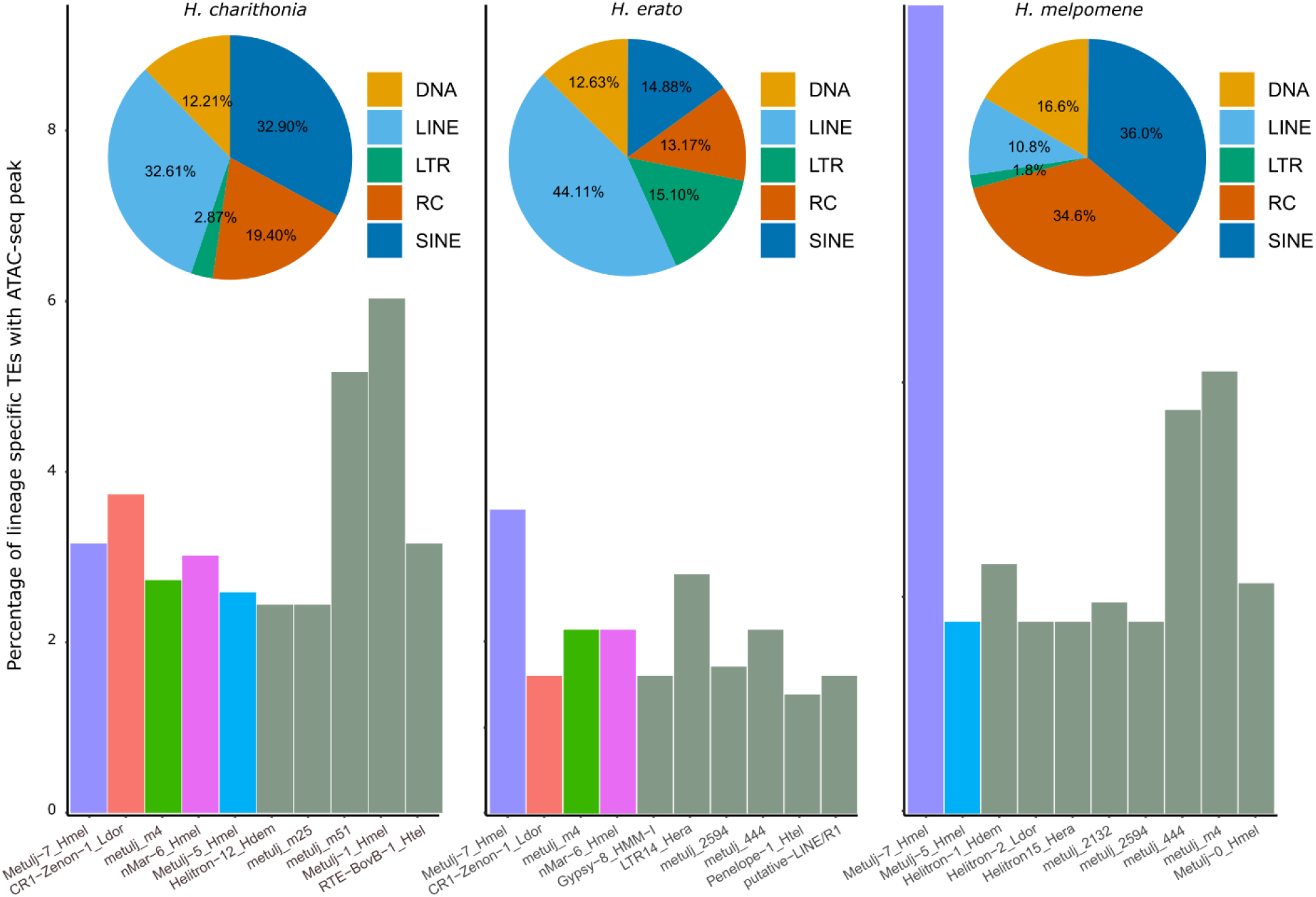
Transposable Element (TE) composition of lineage-specific (i.e., compared to all other species) ATAC-seq peaks within TEs. Pie charts show the TE family distribution. Bar plots show the percentage of the ten most represented TEs in each lineage relative to the total number lineage-specific ATAC-seq peaks within TEs.

## References

Ashraf S, Hu X, Roote J, Ip Y. 1999. The mesoderm determinant Snail collaborates with related zinc-finger proteins tocontrol Drosophila neurogenesis. EMBO J 18: 6426–6438.

Ashraf SI, Ganguly A, Roote J, Ip YT. 2004. Worniu, a snail family zinc-finger protein, is required for brain development in Drosophila. Dev Dyn 231: 379–386.

Bailly-Bechet M, Haudry A, Lerat E. 2014. “One code to find them all”: A perl tool to conveniently parse RepeatMasker output files. Mob DNA 5: 1–15.

Baxter SW, Nadeau NJ, Maroja LS, Wilkinson P, Counterman B a, Dawson A, Beltran M, Perez-Espona S, Chamberlain N, Ferguson L, et al. 2010. Genomic hotspots for adaptation: the population genetics of Müllerian mimicry in the Heliconius melpomene clade. PLoS Genet 6: e1000794.

Bolger AM, Lohse M, Usadel B. 2014. Trimmomatic: A flexible trimmer for Illumina sequence data. Bioinformatics 30: 2114–2120.

Bourque G, Burns KH, Gehring M, Gorbunova V, Seluanov A, Hammell M, Imbeault M, Izsvák Z, Levin HL, Macfarlan TS, et al. 2018. Ten things you should know about transposable elements. Genome Biol 19: 1–12.

Branco MR, Chuong EB. 2020. Crossroads between transposons and gene regulation. Philos Trans R Soc B Biol Sci 375: 2–5.

Buenrostro J, Giresi P, Zaba L, Chang H, Greenleaf W. 2013. Transposition of native chromatin for multimodal regulatory analysis and personal epigenomics. Nat Methods 10: 1213–1218.

Campos JL, Charlesworth B. 2019. The effects on neutral variability of recurrent selective sweeps and background selection. Genetics 212: 287–303.

Cerbin S, Jiang N. 2018. Duplication of host genes by transposable elements. Curr Opin Genet Dev 49: 63–69.

Charlesworth B. 2009. Effective population size and patterns of molecular evolution and variation. Nat Rev Genet 10: 195–205.

Charlesworth B. 2001. The effect of life-history and mode of inheritance on neutral genetic variability. Genet Res 77: 153–166.

Cicconardi F, Lewis JJ, Martin SH, Reed RD, Danko CG, Montgomery SH. 2021. Chromosome Fusion Affects Genetic Diversity and Evolutionary Turnover of Functional Loci but Consistently Depends on Chromosome Size. Mol Biol Evol 38: 4449–4462.

Collins RL, Brand H, Karczewski KJ, Zhao X, Alföldi J, Francioli LC, Khera A V., Lowther C, Gauthier LD, Wang H, et al. 2020. A structural variation reference for medical and population genetics. Nature 581: 444–451.

Conesa A, Götz S, García-Gómez JM, Terol J, Talón M, Robles M. 2005. Blast2GO: A universal tool for annotation, visualization and analysis in functional genomics research. Bioinformatics 21: 3674–3676.

Cutter AD, Payseur BA. 2013. Genomic signatures of selection at linked sites: Unifying the disparity among species. Nat Rev Genet 14: 262–274.

Darling AE, Mau B, Perna NT. 2010. Progressivemauve: Multiple genome alignment with gene gain, loss and rearrangement. PLoS One 5: e11147.

Davey JW, Barker SL, Rastas PM, Pinharanda A, Martin SH, Durbin R, McMillan WO, Merrill RM, Jiggins CD. 2017. No evidence for maintenance of a sympatric Heliconius species barrier by chromosomal inversions. Evol Lett 1: 138–154.

Davey JW, Chouteau M, Barker SL, Maroja L, Baxter SW, Simpson F, Merrill RM, Joron M, Mallet J, Dasmahapatra KK, et al. 2016. Major improvements to the Heliconius melpomene genome assembly used to confirm 10 chromosome fusion events in 6 million years of butterfly evolution. G3 Genes|Genomes|Genetics 6: 695–708.

Davies N, Bermingham E. 2002. The historical biogeography of two Caribbean butterflies (Lepidoptera: Heliconiidae) as inferred from genetic variation at multiple loci. Evolution (N Y) 56: 573–589.

Degiorgio M, Huber CD, Hubisz MJ, Hellmann I, Nielsen R. 2016. SweepFinder2: Increased sensitivity, robustness and flexibility. Bioinformatics 32: 1895–1897.

Diehl AG, Ouyang N, Boyle AP. 2020. Transposable elements contribute to cell and species-specific chromatin looping and gene regulation in mammalian genomes. Nat Commun 11: 1–18.

Ebert P, Audano PA, Zhu Q, Rodriguez-Martin B, Porubsky D, Bonder MJ, Sulovari A, Ebler J, Zhou W, Mari RS, et al. 2021. Haplotype-resolved diverse human genomes and integrated analysis of structural variation. Science (80-) 372.

Edelman NB, Frandsen PB, Miyagi M, Clavijo B, Davey J, Dikow R, García-accinelli G, Van Belleghem SM, Patterson N, Daniel E, et al. 2019. Genomic architecture and introgression shape a butterfly radiation. Science (80-) 366: 24174–24183.

Flynn JM, Hubley R, Goubert C, Rosen J, Clark AG, Feschotte C, Smit AF. 2020. RepeatModeler2 for automated genomic discovery of transposable element families. Proc Natl Acad Sci U S A 117: 9451–9457.

Fueyo R, Judd J, Feschotte C, Wysocka J. 2022. Roles of transposable elements in the regulation of mammalian transcription. Nat Rev Mol Cell Biol 24: 19–24.

Garcia-Perez JL, Widmann TJ, Adams IR. 2016. The impact of transposable elements on mammalian development. Dev 143: 4101–4114.

Guo X, Yin C, Yang F, Zhang Y, Huang H, Wang J, Deng B, Cai T, Rao Y, Xi R. 2019. The cellular diversity and transcription factor code of Drosophila enteroendocrine cells. Cell Rep 29: 4172-4185.e5.

Gupta S, Stamatoyannopoulos JA, Bailey TL, Noble WS. 2007. Quantifying similarity between motifs. Genome Biol 8.

Heinz S, Benner C, Spann N, Bertolino E, Lin YC, Laslo P, Cheng JX, Murre C, Singh H, Glass CK. 2010. Simple combinations of lineage-determining transcription factors prime cis-regulatory elements required for macrophage and B cell identities. Mol Cell 38: 576–589.

Hendrickx F, Corte Z De, Sonet G, Belleghem SM Van, Köstlbacher S, Vangestel C. 2022. A masculinizing supergene underlies an exaggerated male reproductive morph in a spider. Nat Ecol Evol 6: 195–206.

Hill WG, Robertson A. 1966. The effect of linkage on limits to artificial selection. Genet Res (Camb) 8: 269–294.

Iyer EPR, Iyer SC, Sullivan L, Wang D, Meduri R, Graybeal LL, Cox DN. 2013. Functional genomic Analyses of two morphologically distinct classes of Drosophila sensory neurons: Post-mitotic roles of transcription factors in dendritic patterning. PLoS One 8.

Jackman SD, Coombe L, Chu J, Warren RL, Vandervalk BP, Yeo S, Xue Z, Mohamadi H, Bohlmann J, Jones SJM, et al. 2018. Tigmint: Correcting assembly errors using linked reads from large molecules. BMC Bioinformatics 19: 1–10.

Jandrasits C, Dabrowski PW, Fuchs S, Renard BY. 2018. Seq-seq-pan: Building a computational pan-genome data structure on whole genome alignment. BMC Genomics 19: 1–12.

Jay P, Chouteau M, Whibley A, Bastide H, Parrinello H, Llaurens V, Joron M. 2021. Mutation load at a mimicry supergene sheds new light on the evolution of inversion polymorphisms. Nat Genet 53: 288–293.

Joly-Lopez Z, Bureau TE. 2018. Exaptation of transposable element coding sequences. Curr Opin Genet Dev 49: 34–42.

Joron M, Frezal L, Jones RT, Chamberlain NL, Lee SF, Haag CR, Whibley A, Becuwe M, Baxter SW, Ferguson L, et al. 2011. Chromosomal rearrangements maintain a polymorphic supergene controlling butterfly mimicry. Nature 477: 203–206.

Keightley PD, Pinharanda A, Ness RW, Simpson F, Dasmahapatra KK, Mallet J, Davey JW, Jiggins CD. 2014. Estimation of the spontaneous mutation rate in Heliconius melpomene. Mol Biol Evol 32: 239–243.

Kohany O, Gentles AJ, Hankus L, Jurka J. 2006. Annotation, submission and screening of repetitive elements in Repbase: RepbaseSubmitter and Censor. BMC Bioinformatics 7: 1–7.

Kozak KM, Wahlberg N, Neild AFE, Dasmahapatra KK, Mallet J, Jiggins CD. 2015. Multilocus species trees show the recent adaptive radiation of the mimetic Heliconius butterflies. Syst Biol 64: 505–524.

Kozlov A, Jaumouillé E, Almeida PM, Koch R, Rodriguez J, Abruzzi KC, Nagoshi E. 2017. A screening of UNF targets identifies Rnb, a novel regulator of Drosophila circadian rhythms. J Neurosci 37: 6673–6685.

Langmead B, Salzberg SL. 2013. Fast gapped-read alignment with Bowtie 2. Nat Methods 9: 357–359.

Lavoie CA, Ii RNP, Novick PA, Counterman BA, Ray DA. 2013. Transposable element evolution in Heliconius suggests genome diversity within Lepidoptera. Mob DNA 4: 1–10.

Leffler EM, Bullaughey K, Matute DR, Meyer WK, Ségurel L, Venkat A, Andolfatto P, Przeworski M. 2012. Revisiting an old riddle: What determines genetic diversity levels within species? PLoS Biol 10: e1001388.

Lewis JJ, Geltman RC, Pollak PC, Rondem KE, Belleghem SM Van. 2019. Parallel evolution of ancient, pleiotropic enhancers underlies butterfly wing pattern mimicry. Proc Natl Acad Sci 116: 24174–24183.

Lewis JJ, Reed RD. 2019. Genome-wide regulatory adaptation shapes population-level genomic landscapes in Heliconius ed. P. Wittkopp. Mol Biol Evol 36: 159–173.

Lewis JJ, Van Belleghem SM, Papa R, Danko CG, Reed RD. 2020. Many functionally connected loci foster adaptive diversification along a neotropical hybrid zone. Sci Adv 6: 1–11.

Li H, Durbin R. 2010. Fast and accurate long-read alignment with Burrows-Wheeler transform. Bioinformatics 26: 589–595.

Li H, Handsaker B, Wysoker A, Fennell T, Ruan J, Homer N, Marth G, Abecasis G, Durbin R. 2009. The Sequence Alignment/Map format and SAMtools. Bioinformatics 25: 2078–2079.

Liu Y, Ramos-Womack M, Han C, Reilly P, Brackett KLR, Rogers W, Williams TM, Andolfatto P, Stern DL, Rebeiz M. 2019. Changes throughout a genetic network Mask the contribution of Hox gene evolution. Curr Biol 29: 2157-2166.e6.

Livraghi L, Hanly J, Van Belleghem S, Montejo-Kovacevich G, van der Heijden E, Loh LS, Ren A, Warren I, Lewis J, Concha C, et al. 2021. Cortex cis-regulatory switches establish scale colour identity and pattern diversity in Heliconius. Elife 10: e68549.

Lucek K, Gompert Z, Nosil P. 2019. The role of structural genomic variants in population differentiation and ecotype formation in Timema cristinae walking sticks. Mol Ecol 28: 1224–1237.

Mallarino R, Henegar C, Mirasierra M, Manceau M, Schradin C, Vallejo M, Beronja S, Barsh GS, Hoekstra HE. 2016. Developmental mechanisms of stripe patterns in rodents. Nature 539: 518–523.

Martin SH, Davey JW, Salazar C, Jiggins CD. 2019. Recombination rate variation shapes barriers to introgression across butterfly genomes. PLOS Biol 17: e2006288.

Martin SH, Eriksson A, Kozak KM, Manica A, Jiggins CD. 2015. Speciation in Heliconius Butterflies : Minimal Contact Followed by Millions of Generations of Hybridisation. BioRxiv 1–24.

Matschiner M, Barth JMI, Tørresen OK, Star B, Baalsrud HT, Brieuc MSO, Pampoulie C, Bradbury I, Jakobsen KS, Jentoft S. 2022. Supergene origin and maintenance in Atlantic cod. Nat Ecol Evol.

Mérot C, Oomen RA, Tigano A, Wellenreuther M. 2020. A roadmap for understanding the evolutionary significance of structural genomic variation. Trends Ecol Evol 35: 561–572.

Mikkelsen TS, Hillier LW, Eichler EE, Zody MC, Jaffe DB, Yang SP, Enard W, Hellmann I, Lindblad-Toh K, Altheide TK, et al. 2005. Initial sequence of the chimpanzee genome and comparison with the human genome. Nature 437: 69–87.

Moest M, Van Belleghem SM, James J, Salazar C, Martin S, Barker S, Moreira G, Mérot C, Joron M, Nadeau N, et al. 2020. Selective sweeps on novel and introgressed variation shape mimicry loci in a butterfly adaptive radiation. PLoS Biol 18: e3000597.

Montgomery S, Rossi M, Wo M, Merrill R. 2021. Neural divergence and hybrid disruption between ecologically isolated Heliconius butterflies. Proc Natl Acad Sci 118: e2015102118.

Montgomery SH, Merrill RM. 2017. Divergence in brain composition during the early stages of ecological specialization in Heliconius butterflies. J Evol Biol 30: 571–582.

Nishikawa H, Iijima T, Kajitani R, Yamaguchi J, Ando T, Suzuki Y, Sugano S, Fujiyama A, Kosugi S, Hirakawa H, et al. 2015. A genetic mechanism for female-limited Batesian mimicry in Papilio butterfly. Nat Genet 47: 405–409.

Ohtani H, Iwasaki Y. 2021. Rewiring of chromatin state and gene expression by transposable elements. Dev Growth Differ.

Okamoto H, Hirochika H. 2001. Silencing of transposable elements in plants. Trends Plant Sci 6: 527–534.

Ou S, Jiang N. 2018. LTR_retriever: A highly accurate and sensitive program for identification of long terminal repeat retrotransposons.

Pastore M. 2018. Overlapping: a R package for Estimating Overlapping in Empirical Distributions. J Open Source Softw 3: 1023.

Pinharanda A, Martin SH, Barker SL, Davey JW, Jiggins CD. 2017. The comparative landscape of duplications in Heliconius melpomene and Heliconius cydno. Heredity (Edinb) 118: 78–87.

Poelstra JW, Vijay N, Hoeppner MP, Wolf JBW. 2015. Transcriptomics of colour patterning and coloration shifts in crows. Mol Ecol 24: 4617–4628.

Pontis J, Planet E, Offner S, Turelli P, Duc J, Coudray A, Theunissen TW, Jaenisch R, Trono D. 2019. Hominoid-specific transposable elements and KZFPs facilitate human embryonic genome activation and control transcription in naive human ESCs. Cell Stem Cell 24: 724-735.e5.

Quinlan AR, Hall IM. 2010. BEDTools: a flexible suite of utilities for comparing genomic features. Bioinformatics 26: 841–2.

Ray DA, Grimshaw JR, Halsey MK, Korstian JM, Osmanski AB, Sullivan KAM, Wolf KA, Reddy H, Foley N, Stevens RD, et al. 2019. Simultaneous TE Analysis of 19 Heliconiine Butterflies Yields Novel Insights into Rapid TE-Based Genome Diversification and Multiple SINE Births and Deaths. Genome Biol Evol 11: 2162–2177.

Reed RD, Chen PH, Frederik Nijhout H. 2007. Cryptic variation in butterfly eyespot development: The importance of sample size in gene expression studies. Evol Dev 9: 2–9.

Rigal M, Mathieu O. 2011. A “mille-feuille” of silencing: Epigenetic control of transposable elements. Biochim Biophys Acta - Gene Regul Mech 1809: 452–458.

Rodriguez-Caro F, Fenner J, Bhardwaj S, Cole J, Benson C, Colombara AM, Papa R, Brown MW, Martin A, Range RC, et al. 2021. Novel doublesex duplication associated with sexually dimorphic development of Dogface butterfly wings. Mol Biol Evol 38: 5021–5033.

Sato A, Tomlinson A. 2007. Dorsal-ventral midline signaling in the developing Drosophila eye. Development 134: 659–667.

Schiffels S, Durbin R. 2014. Inferring human population size and separation history from multiple genome sequences. Nat Genet 46: 919–925.

Simão FA, Waterhouse RM, Ioannidis P, Kriventseva E V. 2015. BUSCO: assessing genome assembly and annotation completeness with single-copy orthologs. Bioinformatics 31: 3210–3212.

Suntsova M V., Buzdin AA. 2020. Differences between human and chimpanzee genomes and their implications in gene expression, protein functions and biochemical properties of the two species. BMC Genomics 21: 1–12.

Tarailo-Graovac M, Chen N. 2009. Using RepeatMasker to identify repetitive elements in genomic sequences. Curr Protoc Bioinforma Chapter 4: Unit 4.10.

Van’t Hof AE. 2016. The industrial melanism mutation in British peppered moths is a transposable element. Nature 534: 102–105.

Van Belleghem SM, Baquero M, Papa R, Salazar C, Mcmillan WO, Counterman BA, Jiggins CD, Martin SH. 2018. Patterns of Z chromosome divergence among Heliconius species highlight the importance of historical demography. Mol Ecol 27: 3852–3872.

Van Belleghem SM, Rastas P, Papanicolaou A, Martin SH, Arias CF, Supple MA, Hanly JJ, Mallet J, Lewis JJ, Hines HM, et al. 2017. Complex modular architecture around a simple toolkit of wing pattern genes. Nat Ecol Evol 1: 52.

Vassetzky NS, Kramerov DA. 2013. SINEBase: A database and tool for SINE analysis. Nucleic Acids Res 41: 83–89.

Villar D, Berthelot C, Aldridge S, Rayner TF, Lukk M, Pignatelli M, Park TJ, Deaville R, Erichsen JT, Jasinska AJ, et al. 2015. Enhancer evolution across 20 mammalian species. Cell 160: 554–566.

Weisenfeld NI, Yin S, Sharpe T, Lau B, Hegarty R, Holmes L, Sogoloff B, Tabbaa D, Williams L, Russ C, et al. 2014. Comprehensive variation discovery in single human genomes. Nat Genet 46: 1350–1355.

Weissensteiner MH, Bunikis I, Catalán A, Francoijs KJ, Knief U, Heim W, Peona V, Pophaly SD, Sedlazeck FJ, Suh A, et al. 2020. Discovery and population genomics of structural variation in a songbird genus. Nat Commun 11: 1–11.

Wellenreuther M, Mérot C, Berdan E, Bernatchez L. 2019. Going beyond SNPs: The role of structural genomic variants in adaptive evolution and species diversification. Mol Ecol 28: 1203–1209.

Westerman EL, Vankuren NW, Massardo D, Buerkle N, Palmer SE, Kronforst MR. 2018. Aristaless controls butterfly wing color variation used in mimicry and mate choice. Curr Biol 28: 1–6.

Wimmer E, Jäckle H C P, Sm C. 2010. A Drosophila homologue of human Sp1 is a head-specific segmentation gene. J Am Chem Soc 132: 2517–2528.

Yan H, Bombarely A, Li S. 2020. DeepTE: A computational method for de novo classification of transposons with convolutional neural network. Bioinformatics 36: 4269–4275.

Zerbino DR, Johnson N, Juettemann T, Wilder SP, Flicek P. 2014. WiggleTools: Parallel processing of large collections of genome-wide datasets for visualization and statistical analysis. Bioinformatics 30: 1008–1009.

Zhang L, Reifová R, Halenková Z, Gompert Z. 2021. How important are structural variants for speciation? Genes (Basel) 12.

Zhang Y, Liu T, Meyer CA, Eeckhoute J, Johnson DS, Bernstein BE, Nussbaum C, Myers RM, Brown M, Li W, et al. 2008. Model-based analysis of ChIP-Seq (MACS). Genome Biol 9.

